# Amyloplast sedimentation repolarizes LAZYs to achieve gravity sensing in plants

**DOI:** 10.1101/2023.04.17.537121

**Authors:** Jiayue Chen, Renbo Yu, Na Li, Zhaoguo Deng, Xinxin Zhang, Yaran Zhao, Chengfu Qu, Yanfang Yuan, Zhexian Pan, Yangyang Zhou, Kunlun Li, Jiajun Wang, Zhiren Chen, Xiaoyi Wang, Xiaolian Wang, Juan Dong, Xing Wang Deng, Haodong Chen

## Abstract

Gravity controls directional growth of plants, and the classical starch-statolith hypothesis proposed more than a century ago postulates that amyloplast sedimentation in specialized cells initiates gravity sensing, but the molecular mechanism remains mysterious. Here, we report that gravistimulation by reorientation triggers the Mitogen-Activated Protein Kinase (MAPK) signaling-mediated phosphorylation of LAZY proteins, the key regulators of gravitropism accumulated more on the lower side of the plasma membrane in columella cells in regular growth *Arabidopsis*. Phosphorylation of LAZY increases its interaction with several TOC proteins on the surface of amyloplasts, facilitating the translocation of LAZY proteins from the plasma membrane to the amyloplasts. Amyloplast sedimentation subsequently guides LAZY to relocate to the new lower side of the plasma membrane in columella cells, where LAZY induces asymmetrical auxin distribution and differential growth. Together, this study provides a molecular interpretation for the starch-statolith hypothesis: the organelle movement-triggered molecular polarity formation.

## INTRODUCTION

Gravity is a key environmental factor acting on all organisms on earth. In plants, gravity usually directs roots to grow downwards (positive gravitropism) and aerial parts to grow upwards (negative gravitropism)^1, 2^, which may control agricultural traits such as drought tolerance^3^, nutrient absorption^4^, and so on. Gravitropism includes three key processes: gravity sensing, signal transduction, and differential growth response^2, 5^. Since the direction and the magnitude of gravity are almost constant on the surface of the earth, gravitropism is regarded as a posture control, triggered by sensing the tilt of organs relative to the direction of gravity vector^5^.

The starch-statolith hypothesis and Cholodny-Went theory are the dogmas for explaining gravity sensing and response, respectively^6^. The starch-statolith hypothesis was proposed one hundred and twenty years ago, in which sedimentation of amyloplasts (starch-filled plastids) in the statocytes was considered to initiate gravity sensing^7–9^. The columella cells in roots and endodermal cells in shoots were demonstrated to be the statocytes for sensing gravity, and genetic disruption or laser ablation of these cells led to complete disruption of the gravitropism^10–12^. Gravity sensing is further divided into susception (physical response) and signal conversion from physical to physiological information^5^. The sedimentation of starchless plastids in *phosphoglucomutase 1* (*pgm1*) mutants was impaired, resulting in a slower gravitropic response in both roots and shoots^13, 14^. In contrast, disruption of actin accelerated amyloplast sedimentation, resulting in promoted gravitropism^15^. Generally, amyloplast sedimentation was considered to be the susception step of gravity sensing^5^, but the molecular roles of amyloplast sedimentation and the underlying mechanisms of signal conversion were unclear. The Cholodny-Went theory proposed that growth curvature is due to an unequal distribution of auxin between the two sides of the curving organ in plants^16^, and auxin efflux transporter PIN proteins were later proved to contribute to the redirection of auxin and subsequent differential growth in gravitropism^17, 18^.

LAZY family proteins were demonstrated to be critically important for the gravitropic responses of both shoots and roots in many plant species, including, rice^3, 19–21^, maize^22, 23^, *Arabidopsis*^24–26^*, Medicago truncatula*^27^, *Prunus domestica*^25^ and *Lotus japonicus*^28^. High-order *lazy* mutants in *Arabidopsis* showed more exaggerated phenotypes than single mutants, though the phenotypes reported by different groups varied^27, 29, 30^. Our previous study showed that light modulates gravitropism by controlling the expression level of *LAZY4*^31^. A recent study showed that *Arabidopsis* LAZY4 (LZY3) proteins displayed polar distribution on plasma membrane of the columella cells in lateral roots, and recruited RCC1-like domain (RLD) proteins from the cytoplasm to the plasma membrane to promote the re-localization of PIN3 and modulate auxin flow^32^. LAZY4 (AtDRO1) has also been reported to show nuclear localization, but its role there is unclear^33^.

*Altered Response to Gravity 1* (*ARG1*) encodes a DnaJ-like protein and its mutation delayed gravitropism in both root and hypocotyl^34^. Mutating *TOC34*, *TOC75*, *TOC120* or *TOC132*, components of the Translocons at the Outer envelope membrane of Chloroplasts (TOC) complexes, significantly enhanced the *arg1* gravitropic defect, even though either *toc132*^Q730Stop^, *toc75*^G658R^, or *arg1* single mutants showed low to no gravitropic phenotypes^35, 36^. The classical function of TOC proteins is to import proteins into the plastids^37–40^, but whether TOC proteins regulate gravitropism by importing unknown signal factors into amyloplasts is unclear^36^.

Mitogen-Activated Protein Kinase (MAPK) cascades are highly conserved in eukaryotic cells. In plants, the MAPK signaling modules regulate many aspects of growth and environmental responses^41^. The MKK4/MKK5-MPK3/MPK6 module regulates stomatal development and patterning^42^, the inflorescence architecture^43^, lateral root development^44^, and so on. The MKK7-MPK6 module positively regulates hypocotyl gravitropism^45^.

Here, we report that gravistimulation triggers the MKK5-MPK3 module to phosphorylate LAZY proteins, which may promote LAZY proteins to enrich on the surface of amyloplasts via interacting with the TOC proteins. Then, amyloplast sedimentation promotes the translocation of LAZY proteins to the new lower side of the plasma membrane in columella cells, leading to asymmetric auxin distribution and ultimately differential growth. Thus, our study reveals the molecular role of amyloplast sedimentation, providing insights into the molecular mechanism of the starch-statolith hypothesis.

## RESULTS

### *LAZY2, 3,* and *4* are requisite for the gravitropic responses in roots

Previous studies showed that simultaneously disrupting *LAZY2* (*NGR1*), *LAZY3* (*NGR3*) and *LAZY4* (*NGR2*) in *Arabidopsis* via combined T-DNA insertions in the Col or Ws ecotypes resulted in various root gravitropism phenotypes, from negative gravitropism to no clear gravitropism, probably due to the complex ecotype backgrounds or growth conditions^27, 30^. To further understand the functions of LAZY family members, we used a CRISPR/Cas9 system to generate *Arabidopsis lazy2 lazy3 lazy4* (*lazy234*) triple mutants in the Col background. Three lines with different mutated alleles in the three *LAZY* genes were established for phenotypic analyses, designated *lazy234-1*, *lazy234-2* and *lazy234-3* (Figure S1A). With regards to gravitropism growth, either under white light or in the dark, primary roots of wild-type plants always grew towards the gravity vector, whereas those of *lazy234* mutants grew randomly in all directions (Figures 1A and 1B), so did the lateral roots in *lazy234* (Figures S1B and S1C). With regards to the amyloplast sedimentation after reorientation, similar behaviors were observed in wild-type and *lazy234* mutants (Figures S1D and S1E), indicating that these LAZYs are dispensable for amyloplast sedimentation. As the asymmetric distribution of auxin promotes root gravitropism^46^, and *DR5rev::GFP* is widely used to indicate auxin distribution^47^, we mutated *LAZY2, LAZY3,* and *LAZY4* in the *DR5rev::GFP* reporter line, resulting in losing gravitropic responses (Figure S2A-S2D). After the gravistimulation via reorientation, *DR5rev::GFP* signals completely lost their differential distribution in *DR5rev::GFP/lazy234* plants compared with wild-type (Figures 1C, 1D, S2E and S2F). Thus, these LAZY proteins are essential for gravity-triggered polar auxin distribution and bending responses in both primary and lateral roots.

**Figure 1.**
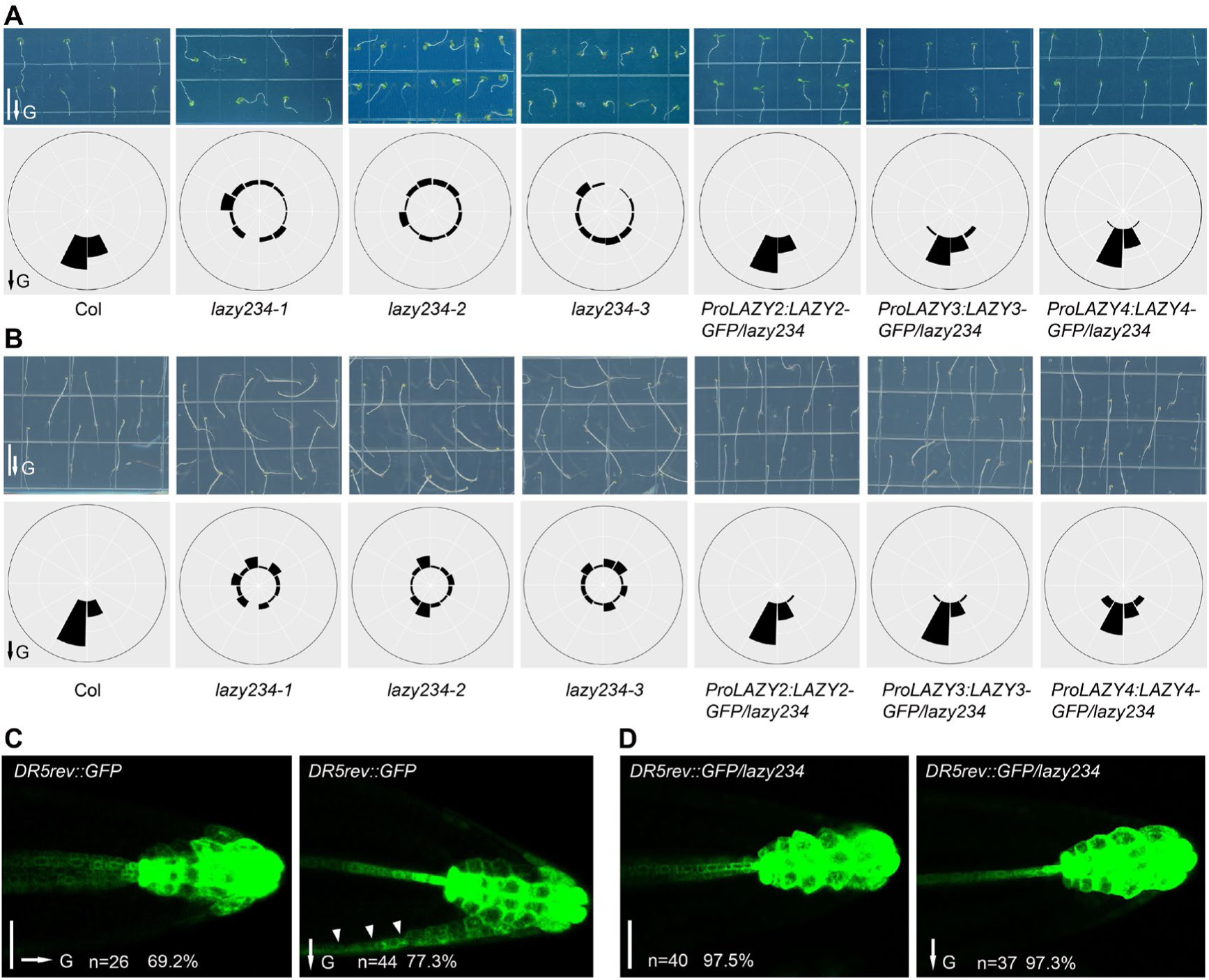
*Arabidopsis* lacking *LAZY2, 3* and *4* completely lose the gravitropic response and gravistimulation-induced auxin redistribution in roots. **(A)** Col, *lazy234* triple mutant, and complementation lines were grown vertically under white light for 4 days. Upper, representative seedlings. Lower, frequencies of root tip angles in each 30° division around a circle (n≥34 for each genotype). Scale bar, 1 cm. **(B)** Col, *lazy234* triple mutant, and complementation lines were grown vertically in the dark for 4 days. Upper, representative seedlings. Lower, frequencies of root tip angles in each 30° division around a circle (n≥26 for each genotype). Scale bar, 1 cm. (**C** and **D**) Expression of *DR5rev::GFP* in wild-type and *lazy234* primary roots before (left) and after gravistimulation via 90° reorientation (right) for around 4 hours. Random primary roots of *DR5rev::GFP/ lazy234* were aligned to the gravity vector before the reorientation, therefore no gravity vector was marked in D, left. Scale bars, 50 μm. Arrowheads indicate the asymmetric accumulation of auxin on one side. In A to D, arrows labeled “G” indicate the direction of gravity. See also Figures S1, and S2.

### LAZY2, 3 and 4 proteins are strongly localized on the amyloplasts and plasma membrane in columella cells

To study protein subcellular localization, the endogenous promoter-driven LAZY-GFP transgenic lines were generated in the *lazy234* mutant background. Each of the transgenes similarly and efficiently rescued the gravitropic phenotypes of *lazy234* primary roots (Figures 1A and 1B). In roots, the three LAZY proteins were predominantly expressed in columella cells, and LAZY4 was also expressed in the endodermis (Figures 2A-2C, and S3A), consistent with the previous results of promoter-driving GUS expression^30^. More strikingly, these LAZY proteins were not only localized on the plasma membranes but also to the surface of the amyloplasts in the columella cells (Figures 2D-2F), which has never been observed before. Interestingly, for LAZY proteins localized on the plasma membrane, they are preferentially accumulated on the plasma membrane regions adjacent to amyloplasts (Figures 2D-2F). Since starch-statolith hypothesis proposed that amyloplast sedimentation initiates gravity sensing, the localization of LAZY proteins on the surface of amyloplasts suggested that they may play critical roles in an early stage of gravity sensing and signal transduction.

**Figure 2.**
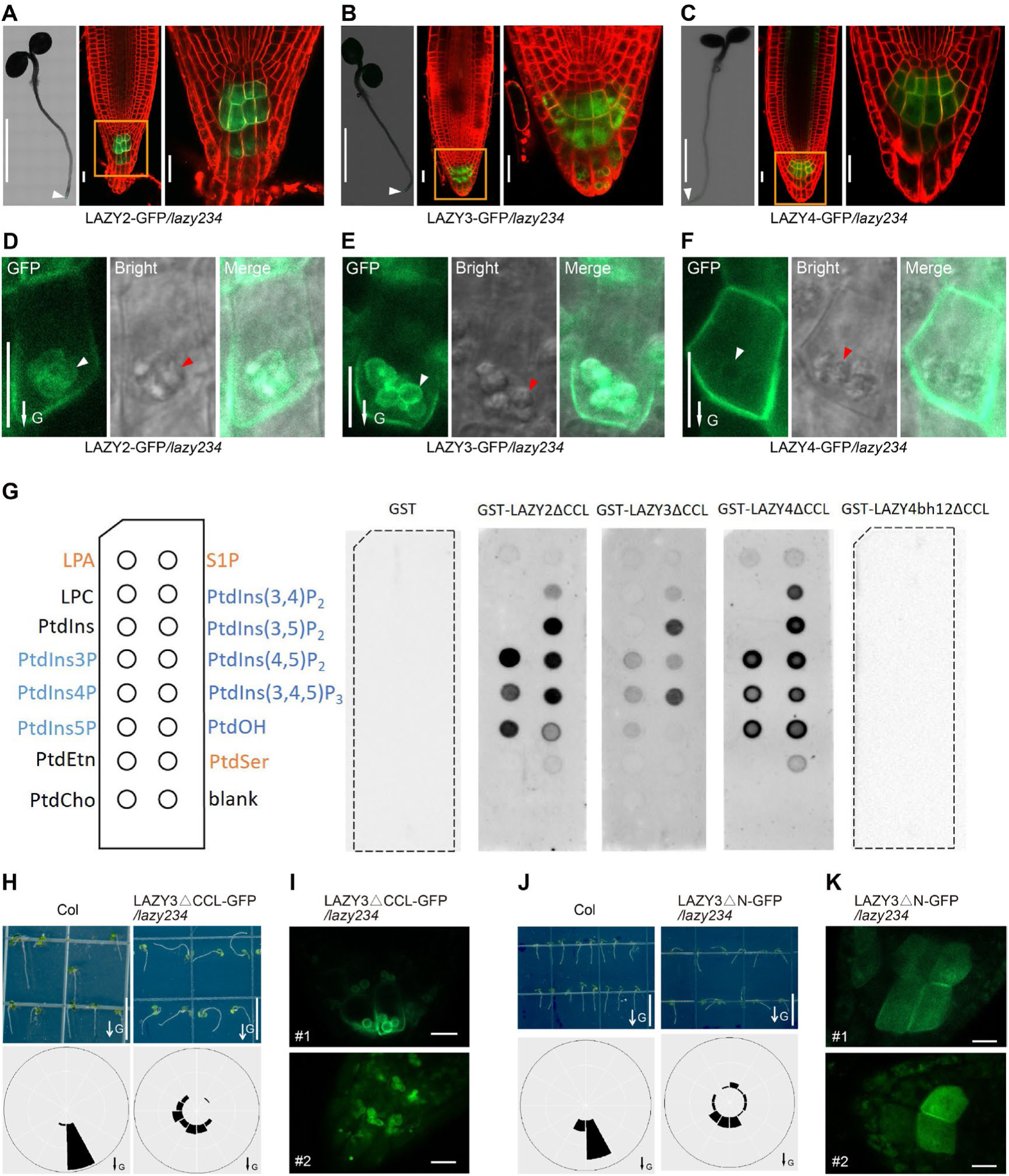
*Arabidopsis* LAZY2, 3 and 4 proteins are predominantly localized on the amyloplasts and plasma membrane of columella cells in roots. **(A to C)** Localization of GFP-tagged LAZY2, LAZY3, and LAZY4 proteins (green) at the root tip of seedlings grown in white light. For each figure panel, left, a confocal image (assembled) shows the predominant expression of LAZY-GFP at the root tip (white arrowhead); middle, a confocal image shows the subcellular localization of LAZY-GFP with an enlarged view (right). Cell outlines were stained with 5 μM FM4-64 for 2 minutes (red). Scale bars, 0.2 cm (left); 20 μm (middle and right). (**D** to **F**) Detailed localization of LAZY2/3/4-GFP protein (green) at the plasma membrane and around the amyloplast envelope in columella cells. White and red arrowheads point to the amyloplasts observed under the Confocal or DIC microscopy, respectively. Scale bars, 10 μm. **(G)** Lipid overlay assay with purified GST, GST-LAZY2ΔCCL, GST-LAZY3ΔCCL, GST-LAZY4ΔCCL and LAZY4bh12ΔCCL using PBS buffer. LPA, lysophosphatidic acid; S1P, sphingosine-1-phosphate; LPC, lysophosphatidylcholine; PtdIns, phosphatidylinositol; PtdIns(x)P_(y)_, mono/bis/tris phosphates, x is the phosphorylation position while y is the number of the phosphate groups; PtdEtn, phosphatidyl-ethanolamine; PtdCho, phosphatidylcholine; PtdOH, phosphatidic acid; PtdSer, phosphatidylserine. LAZY4bh12 means mutating K/R to A within two BH peaks (details in Figure S3C). ΔCCL means the deletion of the last 14 amino acids. **(H)** Col and LAZY3ΔCCL-GFP*/lazy234* transgenic lines were grown vertically under white light for 3 days. Upper, representative seedlings. Lower, frequencies of root tip angles in each 30° division around a circle (n≥45). Scale bars, 1 cm. **(I)** Localization of LAZY3ΔCCL-GFP proteins (green) at the root tip of seedlings grown in white light. Scale bars, 10 μm. **(J)** Col and LAZY3ΔN-GFP*/lazy234* transgenic lines were grown vertically under white light for 3 days. ΔN indicates the deletion of 2-27 amino acids. Upper, representative seedlings. Lower, frequencies of root tip angles in each 30° division around a circle (n≥21). Scale bars, 1 cm. **(K)** Localization of LAZY3ΔN-GFP proteins (green) at the root tip of seedlings grown in white light. Scale bars, 10 μm. In D to F, H and J, arrows labeled “G” indicate the direction of gravity. See also Figure S3.

LAZY2, 3 and 4 proteins were localized on the membranes, but they have no transmembrane domain or lipid acylation modification sites that may contribute to membrane association. Previous studies proved that basic and hydrophobic (BH) regions within proteins may promote their association with membranes^48, 49^. BH score analysis showed that each LAZY has potential membrane binding regions (Figure S3B). Lipid overlay assays showed that these LAZY proteins could bind multiple membrane phospholipids, especially phosphatidylinositolphosphates (PIPs) (Figure 2G). Mutating amino acids K/R to A within the BH peaks of LAZY4 reduced the predicted BH score (Figures S3B and S3C), and disrupted its binding to the membrane phospholipids (Figure 2G). These results suggest that these LAZY proteins may associate with membrane via binding phospholipids.

Since previous studies showed that the conserved C terminus in LAZY1 family proteins (CCL) domain was important for the function of LAZYs^29, 32^, and our CRISPR alleles showed that the N-terminal was also important (Figure S1A), we generated the GFP-fused truncated LAZY3 lacking the CCL or N-terminal (containing domain I) region in *lazy234* background. Both complementation lines showed severe gravitropic defects, supporting the importance of these two regions (Figures 2H and 2J). Deletion of CCL domain appears to increase the accumulation of LAZY3 proteins on the amyloplasts relative to the plasma membrane (Figure 2I). In contrast, deletion of the N-terminal region disrupted the amyloplast localization of LAZY3, resulting in increased cytosol location (Figure 2K). These data indicate that proper distribution of LAZY proteins on the amyloplasts and plasma membrane is critical for their function in gravitropism.

### Sedimentation of amyloplasts promotes the gravity-triggered redistribution of LAZY proteins

Since LAZY proteins were preferentially localized on the lower side of the plasma membrane in columella cells (Figures 2D-2F), we studied the redistribution of LAZY proteins under the regulation of gravity in both primary and lateral roots. Results showed that gravistimulation by re-orientating plants 90° could trigger re-polarization of all three LAZY proteins in columella cells of primary roots, with more proteins accumulated on the new lower side than the upper side (Figures 3A-3D, S3D and S3E). Consistently, in the columella cells of lateral roots, these LAZY proteins also displayed polarization and repolarization after re-orientating the seedlings 180° (Figures S3F and S3G).

**Figure 3.**
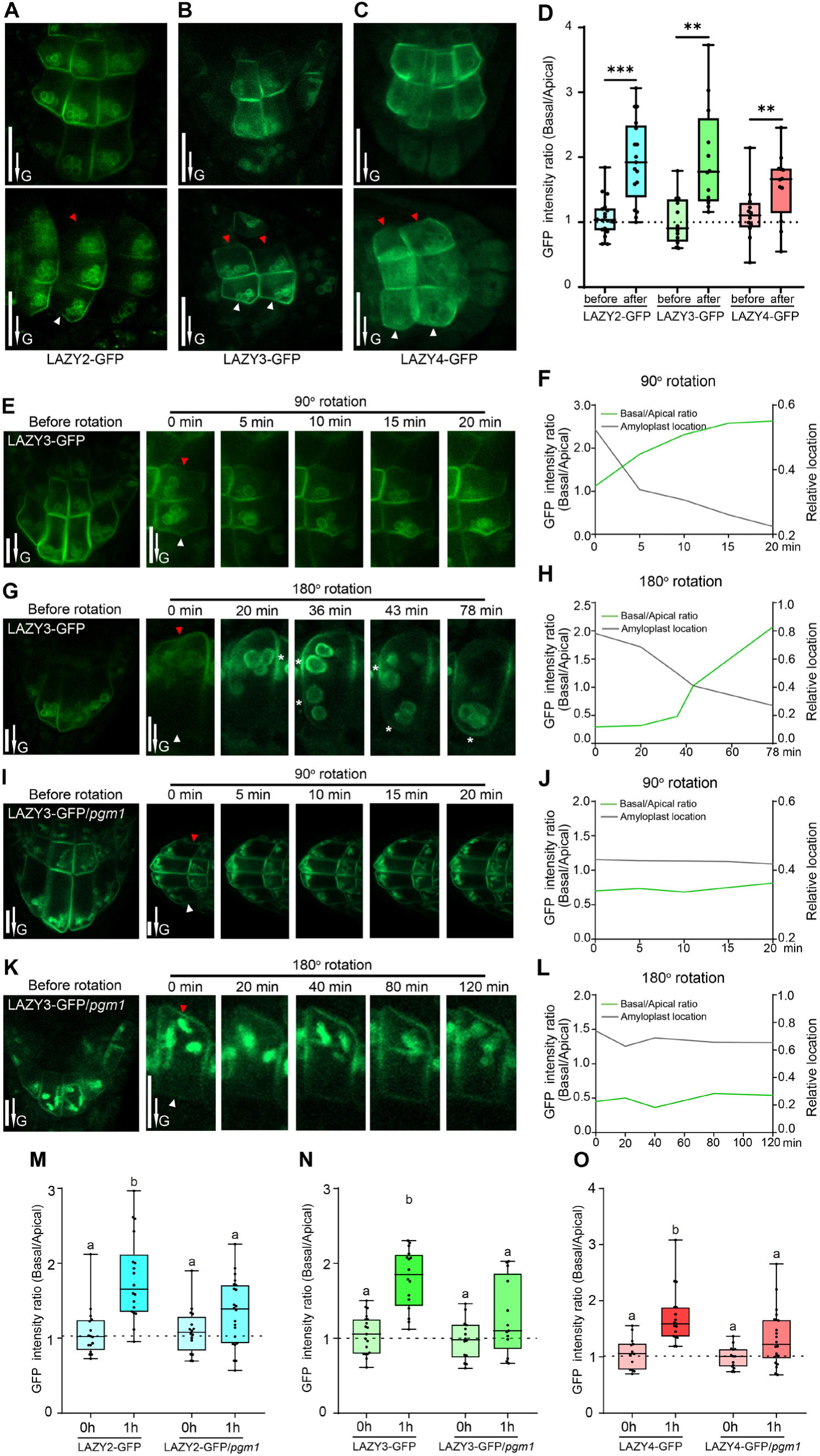
Amyloplast sedimentation promotes the gravity-triggered redistribution of LAZY proteins to the lower side of columella cells. (**A** to **D**) LAZY proteins show polar distribution in columella cells under the control of gravity. (A to C) The seedlings were grown vertically on MS plates, and reoriented 90° and kept for 0.5 to 1 h of gravistimulation. Red and white arrowheads indicate the accumulation of LAZY-GFP proteins on the upper and lower sides of columella cells, respectively. Scale bars, 20 μm. (D) Statistical analysis of fluorescence intensity ratios of the two sides of columella cells (Basal/Apical) in panels A to C. Asterisks indicate Student’s t-test values (***, P < 0.001; **, P < 0.01; n≥12). The method for intensity ratio analysis is shown in Figures S3B and S3C. (**E** to **H**) Gravity-triggered redistribution of LAZY proteins to the lower side of columella cells correlates with the sedimentation of amyloplasts. LAZY3-GFP seedlings were grown vertically and then reoriented 90° (E) or 180° (G), and fluorescence was collected at several time points. Red and white arrowheads indicate the upper and lower sides of columella cells, respectively. White stars in panel G indicate the positions of plasma membrane adjacent to amyloplasts, where the fluorescence is strong. Scale bars, 10 μm. The fluorescence intensity ratio (Basal/Apical) and average relative-locations of the amyloplasts (relative distances from the amyloplasts to the new bottoms of cells, cell heights were set as 1) are shown in F and H. (**I** to **L**) Mutation of *PGM1* delayed both amyloplast sedimentation and redistribution of LAZY3-GFP proteins triggered by gravistimulation. Data collection and analyses were the same as in E to H. (**M** to **O**) Mutation of *PGM1* delayed the redistribution of LAZY2, LAZY3 and LAZY4 proteins. The seedlings were grown vertically and then reoriented 90° and kept 1 h for gravistimulation. The statistical analyses of fluorescence intensity ratios of LAZY-GFP proteins on the two sides of columella cells (Basal/Apical) are shown. One way ANOVA was used to assess the statistical significance (P<0.05; n≥13). Representative images are shown in Figure S4B. In A, B, C, E, G, I and K, arrows labeled “G” indicate the direction of gravity. In E, G, I, and K, the images labeled as “Before rotation” were rotated from the images labeled as “0 min” after 90° or 180° rotation, to show the original orientation. See also Figures S3 and S4.

We further studied whether the repolarization site of LAZY is related to where the amyloplasts sediment in the columella cells. Because LAZY3-GFP showed a strong amyloplast-localized fluorescence signal in our obtained transgenic lines, it was selected as the representative for detailed analysis. After the seedlings were rotated 90° or 180°, we found that redistributed LAZY3-GFP proteins to the new bottom side of the columella cells highly correlated with where the amyloplasts sediment (Figures 3E-3H). Similar coordination was also observed for LAZY2-GFP and LAZY4-GFP (Figure S4A). Furthermore, to corroborate this coordination, we analyzed the distribution of LAZY proteins in the *pgm1* mutant. Indeed, mutation of *PGM1* clearly delayed the sedimentation of plastids after 90° or 180° rotation and we observed slower redistribution of LAZY proteins to the new lower side of the columella cells (Figures 3I-3L). Further statistical analysis clearly showed that gravistimulation-induced polar redistribution of LAZY2, 3, and 4 were all delayed in the *pgm1* mutant (Figures 3M-3O, and S4B). These results suggest that the sedimentation of amyloplasts promote the redistribution of LAZY proteins after gravistimulation.

### Gravistimulation induces phosphorylation of LAZY4 via MKK5-MPK3

To study how gravistimulation mediates the translocation of LAZY proteins, LAZY4-GFP was used as the representative for Immunoprecipitation-Mass Spectrometry (IP-MS) analyses due to its highest expression level (based on fluorescence) among the three LAZYs. LAZY4-GFP seedlings were grown vertically or treated with gravistimulation (0.5 h, 2 h, or 24 h) for phosphor-proteomic analysis. The mass spectrometric analysis identified 13 phosphorylation sites in total, most of which showed up-regulated phosphorylation levels after 0.5-h or 2-h gravistimulation (Figure S5 and Table S1). Phosphorylated peptide “QEITHRPSISSASSHHPR” containing five phosphorylation sites T18/S22/S24/S25/S27 showed the highest abundance based on MS1-intensity-based label-free quantification, and interestingly its phosphorylation levels were induced dramatically by gravistimulation within 0.5 h but reduced to the original status after 24 h (Figure 4A). Furthermore, to identify possible kinases, we searched the IP-MS data of LAZY4-GFP and found the protein abundance of two kinases, MKK5 and MPK3, was increased dramatically after gravistimulation within 0.5 h and decreased to the original level after 24h (Figure 4B), a pattern highly similar to the phosphorylation dynamics of LAZY4 (Figure 4A). Additionally, most of the MPK3 proteins co-immunoprecipitated by LAZY4 are phosphorylated (Figure 4C). Consistently, *in vitro* pull-down experiments also showed that the interactions of LAZY4 with MPK3 and MKK5 were detectable only after the incubation with both kinases (Figure 4D). *In vitro* phosphorylation assay confirmed that MKK5-activated MPK3 can phosphorylate LAZY4, and the phosphorylation sites matched well with the gravistimulation-induced phosphorylation sites (Figure 4E and Table S1). MPK3 and MPK6 are often functionally redundant in regulating developmental processes and the null allele of the double mutant is seedling lethal^42^. To evaluate the root gravitropism phenotypes, we reduced the expression of *MPK3* via inducible RNAi in *mpk6* using the *mpk6^-/-^mpk3RNAi*-est line^44^, and found the double mutants show delayed gravitropic response (Figure 4F). These *in vivo* and *in vitro* data suggest that gravistimulation induces the phosphorylation of LAZY4 by MKK5 and MPK3.

**Figure 4.**
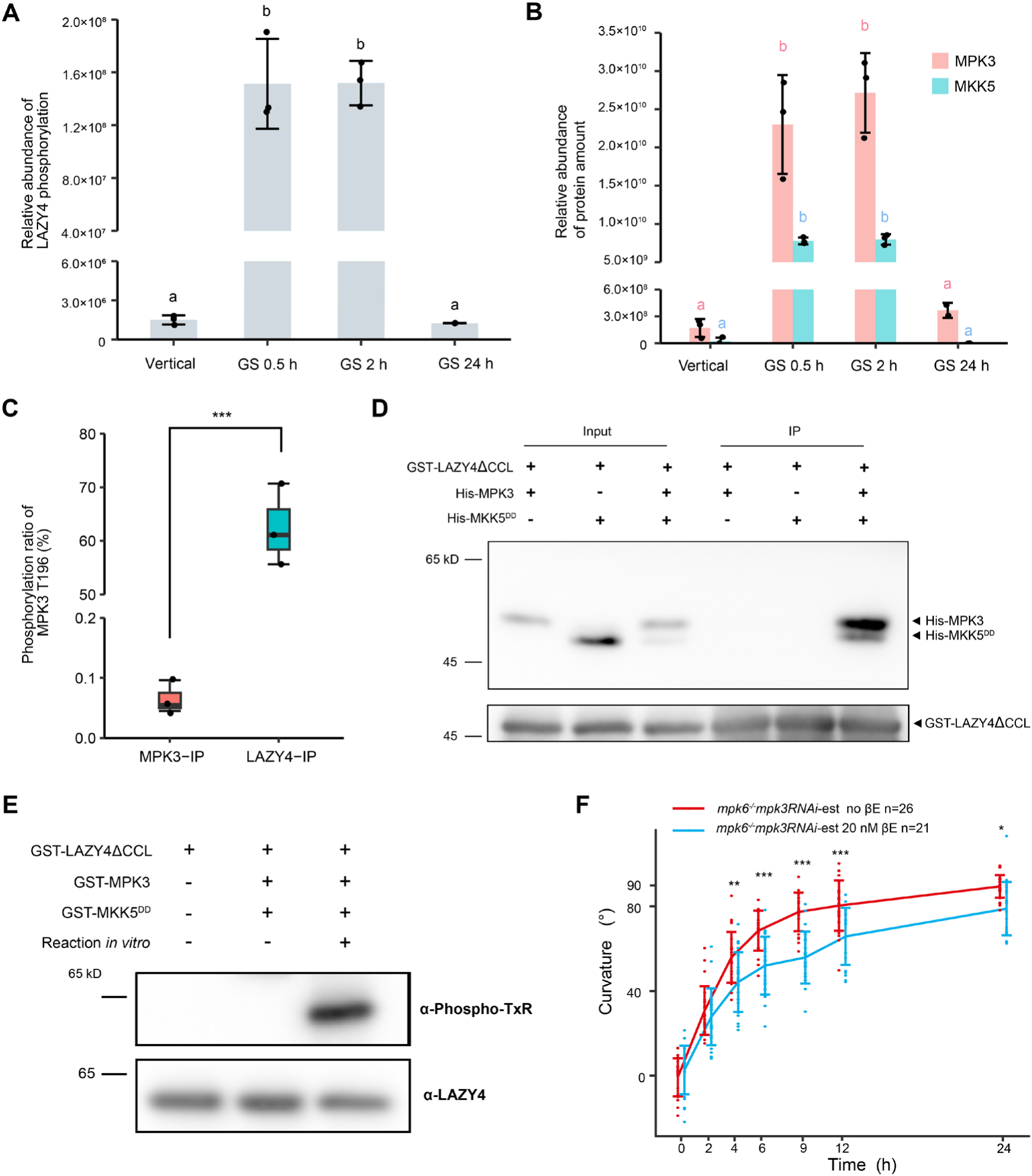
Gravity-induced phosphorylation of LAZY4 by MKK5-MPK3. **(A)** Gravistimulation induces the phosphorylation of LAZY4. LAZY4-GFP seedlings were grown vertically or gravistimulated (90° reorientation) for 0.5 h, 2 h, and 24 h. Then, LAZY4-GFP proteins were immunoprecipitated for Mass-Spectrometry (MS) analysis. The MS result was summarized in table S1, and the relative phosphorylation levels of the representative peptide “QEITHRPSISSASSHHPR” are shown. One way ANOVA was used to assess the statistical significance (p< 0.001; n=3 or 2). **(B)** LAZY4 shows stronger interaction with MPK3 and MKK5 after gravistimulation in *Arabidopsis*. The original data is from the experiment in panel A. The barplot shows the comparison of MPK3 and MKK5 abundance co-immunoprecipitated by LAZY4-GFP. MS1 intensities of all identified peptides of MPK3 and MKK5 in the MS results were summed and normalized using MS1 intensity of bait protein LAZY4-GFP. Abundance of MPK3 and MKK5 in different samples was compared with that in the first group. One way ANOVA was used to assess the statistical significance (p< 0.001; n=3 or 2). **(C)** LAZY4 prefers to interact with phosphorylated MPK3 in *Arabidopsis*. MPK3-GFP seedlings were grown vertically and then gravistimulated (90° reorientation) for 0.5 h. Phosphorylated ratio of the MPK3 peptide containing the T196 site was calculated using the MS data of MPK3-GFP IP samples (MPK3-IP) and LAZY4-GFP Co-IP samples (LAZY4-IP, the original data is from the experiment “GS 0.5 h” in panel A). One way ANOVA was used to assess the statistical significance (p< 0.001; n=3). **(D)** LAZY4 interacts with MPK3 and MKK5 *in vitro*. Recombinant GST-LAZY4ΔCCL, His-MPK3 and His-MKK5^DD^ proteins purified from E.coli were incubated at 4 °C overnight. For the lane 3, His-MPK3 and His-MKK5^DD^ was mixed and treated for 30 min under 37 °C with ATP, and then incubated with GST-LAZY4ΔCCL. Glutathione Sepharose 4B was used for immunoprecipitation. Pellet fractions were separated by SDS-PAGE, and visualized by anti-His and anti-GST immunoblots. GST-LAZY4ΔCCL was used since it is easier to express in E.coli than full length GST-LAZY4. **(E)** MPK3 and MKK5^DD^ phosphorylate LAZY4 *in vitro*. The recombinant proteins were mixed and reacted as indicated. For reaction *in vitro*, recombinant GST-MPK3 and GST-MKK5^DD^ proteins were mixed and treated for 30 min under 37 °C with ATP, and then GST-LAZY4ΔCCL was added and treated for 60 min under the same condition. The proteins were resolved by SDS-PAGE, and visualized by anti-Phospho-TxR (recognizing the phosphorylation of T18 in LAZY4) and anti-LAZY4 immunoblots. The LAZY4 proteins were also analyzed by Mass-Spectrometry, and the phosphorylation sites were summarized in Table S1. **(F)** Mutation of MPK3 and MPK6 delays the gravitropic responses of roots. *mpk6^-/-^ mpk3RNAi*-est mutant line was grown vertically under white light on MS plates containing DMSO (no βE) or 20 nM β-estradiol (20 nM βE) to induce RNAi of *MPK3* expression. Seedlings were grown vertically for 4 days and reoriented 90° for gravistimulation. One way ANOVA was used to assess the statistical significance (***p < 0.001; **p < 0.01; *p<0.05). See also Figure S5 and Table S1.

### Phosphorylation of LAZY4 promotes its interaction with TOC proteins on the surface of amyloplasts

Previous studies showed that mutating genes encoding TOC34, TOC75, TOC120 and TOC132, components of the Translocon of Outer Membrane of Chloroplasts (TOC) complexes, locating on the surface of both chloroplasts and amyloplasts, enhanced the gravitropic defects of *arg1*^35, 36^. Thus, these TOC proteins played positive roles in gravitropism, but the underlying mechanism was unclear. Since LAZY proteins also localize on the surface of amyloplasts (Figures 2 and 3), and the phosphorylation of LAZY4 was induced by gravistimulation (Figure 4), we examined the possible interactions between LAZY4 and TOC proteins and studied whether phosphorylation affected the interactions. Among the 13 *in vivo* phosphorylation sites, we assayed all of them by gradually mutating them to Ala (A) or Asp (D), i.e. LAZY4-A5/D5, the 5 Ser/Thr sites located in the phosphorylated peptide with highest MS1 intensity mutated; LAZY4-A9/D9, the 9 sites identified in our early MS experiments mutated; and LAZY4-A13/D13, all 13 sites mutated (detailed in Table S1). Using the yeast two-hybrid assay, we found that LAZY4 and phosphodead LAZY4 (A5, A9 and A13) did not interact with any TOC proteins (Figures 5A, S6A, S6B and Table S1). In contrast, one phosphomimic LAZY4 variant (LAZY4-D9) interacted strongly with TOC34, TOC120 and TOC132, all of which are receptors on the outer membrane of plastids and usually used for importing non-photosynthetic preproteins (Figures 5A, S6A, S6B and Table S1). Previous study showed that a TOC132 variant without N-terminus that only retains the GTP-binding and membrane domains (TOC132GM) could restore the gravitropic response of *arg1 toc132* to that of *arg1* single mutant^36^. We found that all three phosphomimic LAZY4 variants (LAZY4-D5, LAZY4-D9, and LAZY4-D13) showed strong interaction with TOC132GM (Figures 5A, S6A, S6B, and Table S1). In contrast, the central pore component TOC75 did not interact with these phosphomimic LAZY4 proteins. LAZY4-D9 was selected for localization examination and results showed a high co-localization pattern with TOC132-RFP around the amyloplasts in the columella cells (Figures 5B and 5C). These data suggest that gravistimulation-induced phosphorylation of LAZY4 proteins enhances their interaction with the TOC proteins facing the cytosol, which may promote the translocation of LAZY4 proteins onto the surface of amyloplasts.

**Figure 5.**
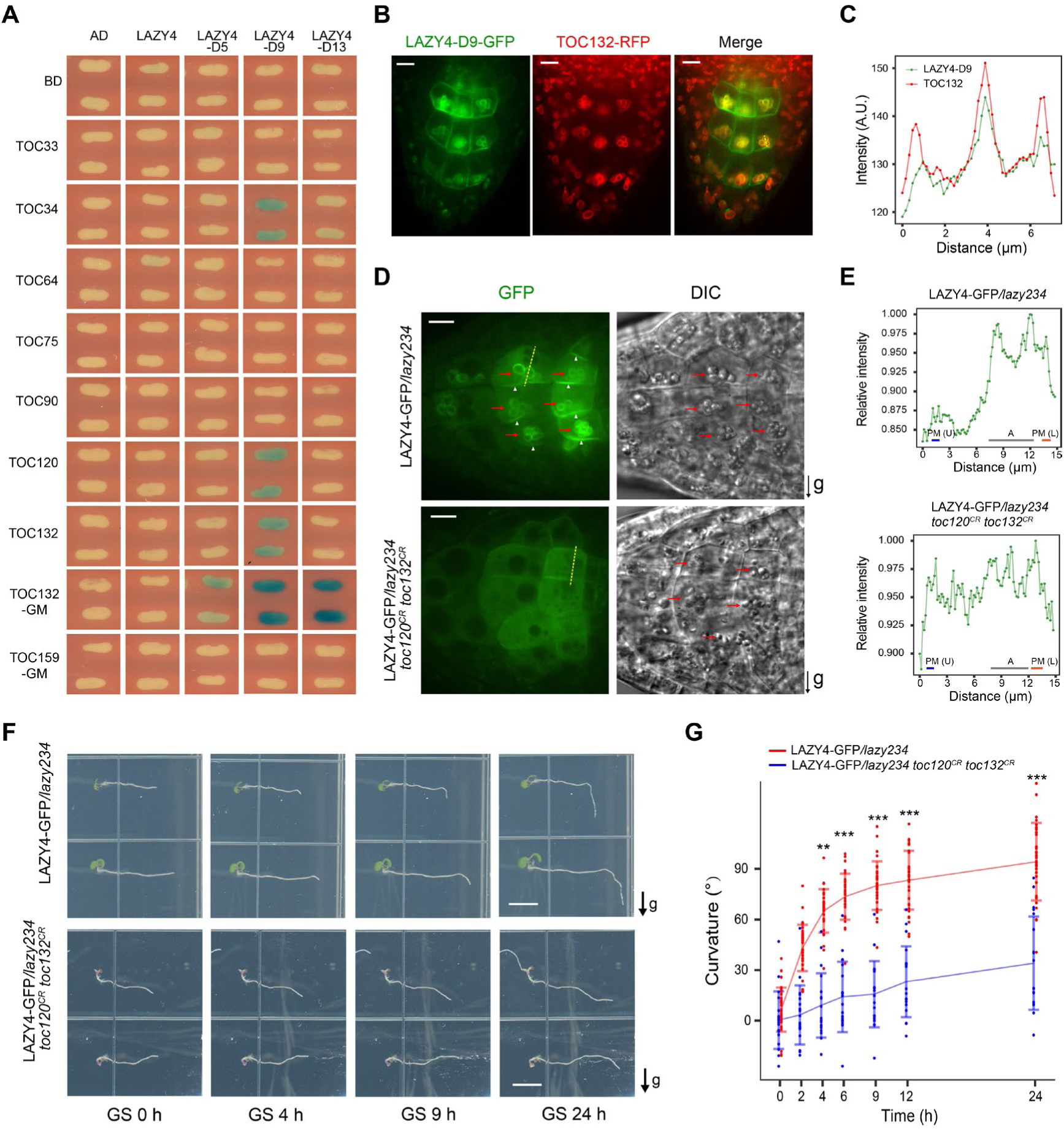
Phosphorylation of LAZY4 promotes its interaction with TOC proteins, which control the redistribution of LAZY4 and the gravitropic response. **(A)** Yeast two-hybrid assays to test interactions between LAZY4 variants and TOC proteins. The phosphorylation sites of LAZY4 were mutated to Asp (D) as indicated in Table S1. Blue indicates positive interactions. Reporter: LacZ; substrate, X-Gal. TOC132GM and TOC159GM indicate the GTP-binding and membrane domains of these TOC proteins. The yeasts were grown for 24 h, and results from 40-h growth with more LAZY4 variants are shown in Figure S6B. **(B** and **C)** Co-localization of LAZY4-D9-GFP and TOC132-RFP in root columella cells. (B) TOC132-RFP was transformed into LAZY4-D9-GFP/*lazy234*, and the seedlings were grown vertically on MS plates for 4 days. Root tip fluorescence was collected. Scale bars, 10 μm. (C) Statistical analysis of fluorescence intensities of LAZY4-D9-GFP and TOC132-RFP in columella cells. Green and red lines show the respective fluorescence intensities of GFP and RFP along the yellow dotted line marked in panel B. a.u., arbitrary units. **(D** and **E)** Mutation of *TOC120* and *TOC132* disrupts the localization of LAZY4-GFP onto the surface of amyloplasts and polar redistribution of LAZY4-GFP on the plasma membrane. (D) LAZY4-GFP/*lazy234* and LAZY4-GFP/*lazy234 toc120*^CR^ *toc132*^CR^ seedlings were grown vertically on MS plates for 4 days. The seedlings were transferred to MS plates with 10 μM CHX to inhibit new protein synthesis, and reoriented 90° for 0.5 to 1-h gravistimulation. Red arrows indicate localization of LAZY4-GFP on the amyloplasts (left) or amyloplasts under DIC (right). White arrowheads indicate the accumulation of LAZY4-GFP proteins on the lower side of columella cells. Scale bars, 10 μm. (E) Statistical analysis of LAZY4-GFP intensity in columella cells. Green lines show the fluorescence intensities of LAZY4-GFP along the yellow dotted line marked in figure D. PM (U), upper side of the plasma membrane; PM (L), lower side of the plasma membrane; A, amyloplasts. **(F** and **G)** Mutation of TOC120 and TOC132 interrupts the gravitropic responses of roots. LAZY4-GFP/*lazy234* and LAZY4-GFP/*lazy234 toc120*^CR^ *toc132*^CR^ seedlings were grown vertically on MS plates for 4 days, and then rotated 90° to the horizontal position for gravistimulation. (F) Representative seedlings. Scale bars, 0.5 cm. (G) Root curvature angle data are means ± SD (n≥20). One-way ANOVA was used to assess the statistical significance (***p < 0.001; **p < 0.01). In D to G, LAZY4-GFP/*lazy234 toc120*^CR^ *toc132*^CR^ seedlings were selected from T2 population based on the cotyledon phenotype (chlorophyll deficiency) and confirmed by sequencing (Table S2). See also Figure S6, Tables S1 and S2.

### TOC proteins are essential for amyloplast localization and polar redistribution of LAZY4 proteins

To analyze the physiological significance of gravistimulation-induced phosphorylation of LAZY4 and its enhanced interaction with TOC proteins, we generated plant reagents expressing mutated TOC proteins or LAZY4 with altered phosphorylation sites. Since LAZY4-D9 showed the strongest interaction with TOC proteins among the LAZY4 variants, we transformed LAZY4-D9-GFP (phosphomimic) and LAZY4-A9-GFP (phosphodead) into *lazy234* mutant. Both transgenic lines appeared to rescue the gravitropic defects of *lazy234* but some LAZY4-A9-GFP lines were statistically less efficient compared to the wild-type LAZY4 in complementation (Figures S6C and S6D). The results indicate that these 9 sites play roles in gravitropism but the effect might be masked by additional phosphorylation sites. Further, we mutated *TOC120* and *TOC132* simultaneously with CRISPR/Cas9 in LAZY4-GFP/*lazy234* since these two *TOC* genes showed redundancy in chlorophyll synthesis (Figure S6A) ^37, 38^. When the plants were gravistimulated (via 90° reorientation) with CHX treatment to inhibit new protein synthesis, mutation of *TOC120* and *TOC132* disrupted the localization of LAZY4-GFP proteins onto the amyloplasts and their polar redistribution on the plasma membrane (Figures 5D and 5E). Consistently, these plants with *toc120* and *toc132* mutations showed strong gravitropic defects (Figures 5F and 5G). In addition, the strong association of LAZY4-D9 with the amyloplast was disrupted by mutations in *TOC120* and *TOC132* (Figures S6E), further supporting that the localization of LAZY4 on amyloplasts is dependent on TOC120 and TOC132. Thus, these TOC proteins are the key anchor proteins on the surface of amyloplasts that bind and facilitate the redistribution of LAZY proteins that are required for asymmetric root growth in plant gravitropic responses.

## DISCUSSION

Gravity sensing, signal transduction, and differential growth are three sequential processes in plant gravitropism^2, 5^. The starch-statolith hypothesis is the most widely-accepted theory explaining gravity sensing and is considered a dogma in the field of plant gravitropism^5–9, 50^, but the underlying molecular effects of amyloplast sedimentation are unknown. Here, we revealed that the molecular effect of amyloplast sedimentation is to promote the redistribution of LAZY proteins via the TOC proteins on the amyloplast surface. Based on this and previous studies, we suggest the following model for gravity sensing in plants (Figure 6): (i) During regularly vertical growth, amyloplasts settle to the bottom in root columella cells, and more LAZY proteins accumulate on the lower side of the plasma membrane. (ii) When plants tilt relative to the direction of gravity vector, gravistimulation induces the interaction between MKK5-MPK3 kinases and LAZY, resulting in phosphorylation of LAZY proteins. (iii) Phosphorylated LAZY proteins show stronger interactions with TOC34/120/132 proteins on the surface of amyloplasts, which may promote their translocation from the plasma membrane to the amyloplasts. (iv) Amyloplast sedimentation brings and guides the LAZY proteins toward the new bottom side of columella cells. In this process, LAZY proteins may keep on trafficking between the amyloplasts and adjacent plasma membrane. When the amyloplasts approach the new bottom, LAZY proteins are accumulated more on the new lower side of the plasma membrane, re-establishing their polarity to promote asymmetric distribution of auxin and bending growth. (v) When the roots resume vertical growth, the LAZY proteins are dephosphorylated by unknown mechanisms, returning to their original state. Although the underlying mechanism of gravistimulation-triggered interaction between MPK3 and LAZY4 still need further investigation, the phosphorylation status of LAZY4 and its interaction with MPK3 can be regarded as two molecular indicators of plant responses to gravistimulation (Figure 4), which must be very useful in future studies in this field. Polarized LAZYs may regulate the asymmetric distribution of auxin through RLD and PIN proteins to mediate differential growth^32^. Since LAZY orthologs are prevalent and have also been demonstrated to regulate gravitropism in many other plant species, including rice^3, 19–21^, maize^22, 23^, *Medicago truncatula*^27^, *Prunus domestica*^25^ and *Lotus japonicus*^28^, and TOC proteins are conserved in plant kingdom^37–40^, it is possible that the mechanism revealed in this study is also applicable to other plants.

**Figure 6.**
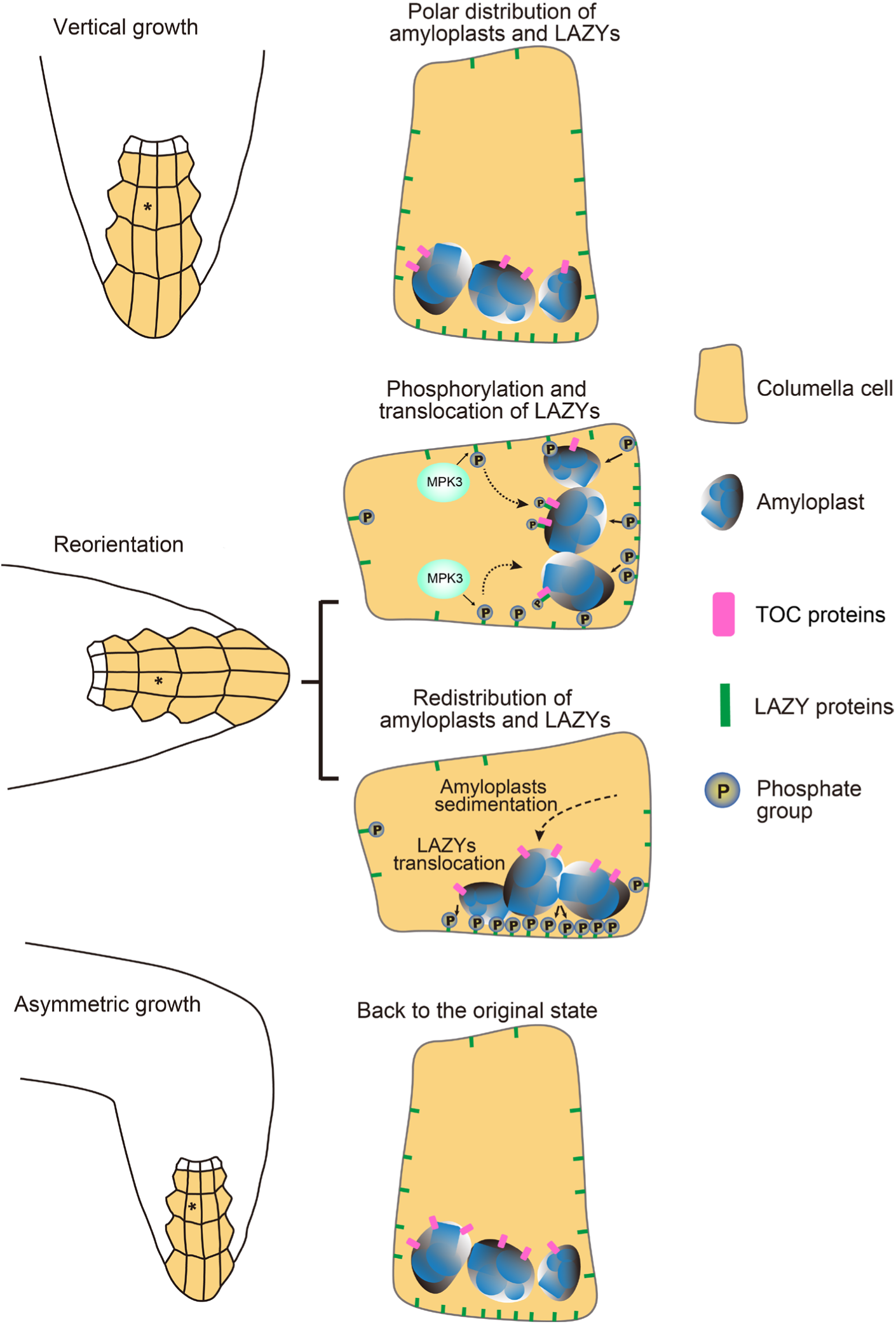
Model for gravity sensing in plant columella cells. Under vertical growth, LAZY proteins are accumulated more on the lower side of the plasma membrane in columella cells. Gravistimulation via reorientation triggers the interactions between the MKK5-MPK3 kinase module and LAZY proteins, resulting in phosphorylation of LAZY proteins. Subsequently, phosphorylated LAZYs may translocate onto the surface of amyloplasts via directly interacting with TOC proteins. Amyloplast sedimentation guides the LAZY proteins to distribute onto the new lower side of the plasma membrane in columella cells, where LAZY induces asymmetrical auxin distribution and differential growth. When the roots resume vertical growth, the LAZY proteins are dephosphorylated, returning to their original state.

For the starch-statolith hypothesis, two models have been proposed to explain how the physical stimulus of amyloplast sedimentation is transformed into a biochemical signal responsible for the gravitropic curvature in plants^51^. The first model postulates that sedimenting statoliths may activate mechano-sensitive ion channels via exerting a pressure on sensitive membranes or cytoskeletons within the statocyte cells^52, 53^, but no such plant mechanosensitive ion channel has been identified. The second model proposes that the contact of statoliths with membrane-bound receptors rather than pressure or tension exerted by the weight of statoliths may achieve graviperception^54^, but the information about the receptor is unavailable. Our study proposes a new model solely involving organelle movement that triggers the repolarization of the key gravitropism regulator LAZYs (Figure 6), suggesting that the molecular effect of amyloplast sedimentation probably doesn’t need mechano-sensitive ion channels or receptors as proposed in the previous models. Interestingly, although the classical function of TOC proteins is to import proteins into plastids^37–40^, it seems that relocation of LAZYs does not require them to be imported into amyloplasts (Figures 2, 3 and 5). This unconventional working mechanism of TOCs makes repolarization of LAZYs simple and fast, resulting in a rapid gravitropic response. Since both movement of organelles and polarity formation are common phenomena, by revealing that the movement of a specific organelle directly triggers the protein re-polarity within cells, our study may also inspire other studies related to polarity.

### Limitations of the study

Although our study provides new insights into the molecular mechanism for the starch-statolith hypothesis, there are a few open issues. First, the underlying mechanism of gravistimulation-triggered interaction between MKK5-MPK3 and LAZY proteins needs further studies. What are the earliest signaling events after gravistimulation and whether MAPKKKs and/or other signaling molecules participate in this process remain the most exciting and challenging questions in the field. Second, we clearly showed that phosphorylation of LAZY4 increased its interaction with TOC proteins, and mutating TOCs interrupted the localization of LAZY4 onto amyloplasts, resulting in severe gravitropic defects. We thus expected that mutating several key phosphorylation sites within LAZY4 would lead to strong gravitropic defects, but this phenomenon only appeared in some transgenic lines under specific growth conditions. It is possible that more phosphorylation sites than we mutated are involved in the interactions between LAZY and TOC proteins *in vivo*. Alternatively, there may exist other uncharacterized mechanisms besides phosphorylation that also contribute to the interaction. Third, the detailed trafficking processes of LAZY proteins between amyloplasts and the plasma membrane need future investigation. For example, how LAZY proteins are unloaded from amyloplasts and how the MAPK-mediated phosphorylation events crosstalk with the RLD-mediated cellular processes to ensure the changes of LAZY polarization in gravitropism.

## STAR METHODS

Detailed methods are provided in the online version of this paper.

## SUPPLEMENTAL INFORMATION

Supplemental information can be found online.

## ACKNOWLEDGMENTS

We thank Prof. Tongda Xu for *mpk6^-/-^mpk3RNAi*-est seeds, Profs. Jian-Kang Zhu and Qijun Chen for CRISPR/Cas9 plasmids, and Profs. Li-Jia Qu, Tongda Xu, William Terzaghi, Genji Qin, Yule Liu, Guangshuo Ou, and Li Yu for helpful comments, Jingxi Sun and Huifang Qin for technical assistance. We thank the National Center for Protein Sciences at Peking University in Beijing, China, particularly Drs. Dong Liu, and Qi Zhang for technical assistance with the mass spectrometry experiments. This study was supported by the National Natural Science Foundation of China (32022005, 31621001, 32170287) and the Tsinghua-Peking Center for Life Sciences. J.D. was funded by grants from the National Institute of Health (GM1851907) and the National Science Foundation (2049642).

## AUTHOR CONTRIBUTIONS

H.C. conceived and supervised the study. Y.Y., R.Y., and K.L. generated *lazy* triple mutants in Col. and *DR5rev::GFP*, and collected the phenotype and fluorescence. R.Y., Y. Zhao, X.Z., J.C., J.W., and X.Y.W. generated the LAZY-GFP lines and collected the fluorescence data. J.C. carried out the lipid overlay assay. Y. Zhao, N.L., and Z.P. generated the LAZY-GFP/*pgm1* lines and collected the phenotype and fluorescence data. J.C., N.L., and C.Q. carried out IP-MS. Y. Zhao, N.L., R.Y., J.C., and Z.P. performed the yeast two hybrid analysis. J.C. and C.Q. performed the *in vitro* phosphorylation and interaction assay. J.C., Y. Zhou, N.L., Z.D., C.Q. and Z.C. generated mutated LAZY4-GFP transgenic lines and collected the phenotype and fluorescence data. X.Z. generated truncated LAZY3-GFP transgenic lines and collected the phenotype and fluorescence data. J.C. and Z.D. analyzed the phenotype of *mpk* mutant. N.L. generated the materials with LAZY4-GFP and LAZY4-D9-GFP in *toc* mutant and collected the phenotype and fluorescence data with the help of X.L.W. Z.D. did multiple analyses about fluorescence, phenotypes, and MS data. J.C., R.Y., N.L., Z.D., Y.Zhao, C.Q., J.D., X.W.D. and H.C. drafted the manuscript.

## DECLARATION OF INTERESTS

The authors declare no competing interests.

**STAR METHODS**

**KEY RESOURCES TABLE**

**RESOURCE AVAILABILITY**

### Lead contact

Further information and requests for resources and reagents should be directed to and will be fulfilled by the lead contact, Haodong Chen

### Materials availability

Constructs and unique reagents generated in this study will be available from the lead contact upon request.

### Data and code availability

All data reported in this paper are available from the lead contact upon request.

## EXPERIMENTAL MODEL AND SUBJECT DETAILS

### ***Arabidopsis* strains**

All wild-type plants used in this study were the Columbia ecotype (Col) of *Arabidopsis thaliana*. Several mutants and transgenic lines used in this study were described previously: *pgm1*^13, 14^, *mpk6^-/-^mpk3RNAi*-est line^44^, and *DR5rev::GFP*^47, 55^.

### ***Arabidopsis* growth**

The seeds were surface-sterilized by soaking in 15% NaClO for around 10 min and washing in sterile distilled water for at least three times. After surface-sterilization, the seeds were plated on MS medium (4.4 g/L MS, 1% sucrose, 0.8-1% agar, pH 5.8), and cold-treated at 4 °C for 2 to 3 days in the dark. The seedlings were grown under white light or dark conditions.

## METHOD DETAILS

### Generation of mutants and transgenic plants

For mutating *Arabidopsis LAZY2*, *LAZY3*, and *LAZY4* simultaneously, we generated a modified Cas9 vector named as pEC-Cas9. First, rbcS-E9 terminator was cloned from pHEE401E^56^, and inserted between the *Bam*H I and *Eco*R I sites of p35S-Cas9-SK^57^ to replace the Nos terminator, resulting in p35S-Cas9-rbcS-E9t. Second, p35S-Cas9-rbcS-E9t was further modified by deleting a *Xho* I site upstream of the double CaMV 35S promoter and adding a *Nhe* I site between *Sal* I and *Hin*d III to generate p35S-Cas9-rbcS-E9t-M. Third, Ec1.2enEc1.1 promoter was cloned from pHEE401E^56^ and inserted between *Nhe* I and *Xho* I sites to generate pEc1.2enEc1.1p-Cas9-rbcS-E9t (*abbr.*: pEC-Cas9). Two sgRNAs were designed for each *LAZY* gene (*LAZY2, LAZY3,* and *LAZY4*), and were inserted into the *Bbs* I sites of the pAtU6-26-M^58^. The primers used for annealing are listed in Table S3. The expression cassettes of six sgRNAs were digested with *Kpn* I and *Sal* I and ligated into pEC-Cas9. Then, the Cas9 cassettes were subcloned into the *Kpn* I and *Eco*R I sites of pCAMBIA1300 (Hyg) and pJIM19 (Gent) vectors to generate pCAMBIA1300-LAZY2/3/4 sgRNA and pJIM19-LAZY2/3/4 sgRNA. These binary constructs were transformed into wild type and *DR5rev::GFP* to generate *lazy2 lazy3 lazy4* (*lazy234*) and *DR5rev::GFP/lazy234* mutant lines. The mutants were analyzed using the primers listed in Table S3, and the lines *lazy234-1*, *lazy234-2*, *lazy234-3*, and *DR5rev::GFP/lazy234* were obtained.

To construct LAZY-GFP transgenic lines driven by the native promoter, the genomic DNAs of *LAZY2*, *LAZY3* and *LAZY4* including promoter regions were amplified by the primers listed in Table S3. Genomic fragments of *LAZY2*, *LAZY3* and *LAZY4* fused with GFP at the C-terminus were inserted into the *Sbf* I*/Xba* I, *Sbf* I*/Xho* I and *Sbf* I*/Spe* I sites of the plant binary vector pJIM19 (Bar) to construct pJIM19-LAZY2-GFP, pJIM19-LAZY3-GFP and pJIM19-LAZY4-GFP. Then, these LAZY-GFP constructs were transformed into wild type (Col) and *lazy234* respectively, and transgenic plants were selected with glufosinate ammonium (20 μg/mL). The obtained plants include *ProLAZY2:LAZY2-GFP* (*abbr.*: LAZY2-GFP/Col, only used in lateral roots fluorescence analysis in Figures S3F and S3G), *ProLAZY2:LAZY2-GFP/lazy234* (*abbr.*: LAZY2-GFP*/lazy234* or LAZY2-GFP), *ProLAZY3:LAZY3-GFP/lazy234* (*abbr.*: LAZY3-GFP*/lazy234* or LAZY3-GFP), and *ProLAZY4:LAZY4-GFP/lazy234* (*abbr.*: LAZY4-GFP*/lazy234* or LAZY4-GFP). These transgenic lines were crossed with *pgm1* to generate LAZY2-GFP*/lazy234/pgm1* (*abbr.*: LAZY2-GFP*/pgm1*), LAZY3-GFP*/lazy234/pgm1* (*abbr.*: LAZY3-GFP*/pgm1*) and LAZY4-GFP*/lazy234/pgm1* (*abbr.*: LAZY4-GFP*/pgm1*). The GFP-fused truncated LAZY3 driven by the *LAZY3* native promoter were amplified using the primers listed in Table S3 and inserted into pJIM19 (Bar). The obtained plants include *ProLAZY3:LAZY3ΔN-GFP/lazy234* (*abbr.*: LAZY3ΔN-GFP/*lazy234)*, and *ProLAZY3:LAZY3ΔCCL-GFP/lazy234* (*abbr.*: LAZY3ΔCCL-GFP/*lazy234*).

To construct *ProLAZY4:LAZY4-D9-GFP/lazy234* and *ProLAZY4:LAZY4-A9-GFP/lazy234* transgenic plants, the promoter and coding sequences of *LAZY4* were amplified from wild type plants using the primers listed in Table S3, and point mutations were generated by overlap PCR. Nine amino acids (T18, S22, S24, S25, S27, S139, S140, S148, S164) were mutated to be Asp (D) or Ala (A) to generate phosphomimic LAZY4-D9 and phosphodead LAZY4-A9, respectively. Then, the fragments were inserted into the *Sbf* I*/Spe* I sites of pJIM19 (Bar), and transformed into *lazy234* triple mutants. Transgenic plants were selected with glufosinate ammonium (20 μg/mL). The obtained plants include *ProLAZY4:LAZY4-D9-GFP/lazy234* (*abbr.*: LAZY4-D9-GFP) and *ProLAZY4:LAZY4-A9-GFP/lazy234* (*abbr.*: LAZY4-A9-GFP).

To construct LAZY4-GFP/*lazy234 toc120*^CR^ *toc132*^CR^ and LAZY4-D9-GFP/*lazy234 toc120*^CR^ *toc132*^CR^, two sgRNAs were designed for each *TOC* (*TOC120* and *TOC132*), and were inserted into the *Bbs* I sites of the pAtU6-26-M^56^ vector. The primers used for annealing are listed in Table S3. The expression cassettes of four sgRNAs were digested with *Kpn* I and *Sal* I and inserted into the *Kpn* I and *Eco*R I sites of pUBQ10:Cas9-P2A-GFP (Gent) ^58^ vector. This construct was transformed into LAZY4-GFP*/lazy234* and LAZY4-D9-GFP/*lazy234* transgenic lines. The mutants were analyzed using the primers listed in Table S3.

To construct TOC132-RFP/LAZY4-D9-GFP/*lazy234* transgenic lines, the genomic DNAs of *TOC132* including promoter regions and coding sequences were amplified by the primers listed in Table S3. Genomic fragments of TOC132 were inserted into the *Sal* I and *Bam*H I sites of the plant binary vector *pCAMBIA1300(Hyg)-mRFP*. Then, the *pCAMBIA1300(Hyg)-TOC132-mRFP* construct was transformed into LAZY4-D9-GFP/*lazy234* transgenic lines, and transgenic plants were selected with Hygromycin B (50 μg/mL).

### Gravitropic phenotype analyses

To study gravitropism of primary roots under regular growth, seedlings were grown 4 days on vertically-oriented MS plates under white light or darkness. To study the tropic responses of primary roots after gravistimulation, seedlings were grown vertically under white light for around 4 days, and then the plates were re-orientated 90°. To study the growth orientations of lateral roots (LR), seedlings were grown vertically on MS plates under white light. Then, the primary roots were aligned to the gravity vector on the 7^th^ day, and the plants continued to grow for several additional days. Root growth angles were measured by Image J software^59^. The distribution frequency of root growth angles was plotted using the R statistics program (http://www.rproject.org).

### Microscopic observation and measurement of signal intensity ratio

For fluorescence observation of *Arabidopsis* seedlings under vertical growth or gravistimulation, the fluorescent images were collected using the confocal laser microscope (Zeiss LSM800 or Andor Dragonfly 200) with a vertical objective table. 488 nm laser was used to detect the fluorescence of LAZY-GFP (including mutated variants), and 561 nm laser was used to detect FM4-64 or TOC132-RFP. Differential interference contrast microscopy (DIC) was used to collect the images of the amyloplasts. Around four-day-old seedlings grown under white light were used for fluorescence observation unless specified otherwise.

Signal intensity was calculated by ImageJ^59^. Signal intensity ratios of LAZY-GFP proteins in columella cells after gravistimulation were calculated by comparing the fluorescence of the lower outer and upper outer lateral plasma membranes of central columella cells or lateral columella cells (the columella cells adjacent to central columella cells). Signal ratio for one root was calculated as an average of signal intensity ratios within this root^60^. The details of signal intensity ratio analysis are shown in Figures S3D and S3E.

### Lipid overlay assay

Recombinant GST-tagged proteins were expressed and purified using BL21 (DE3). Strips with 15 different phospholipids on a nitrocellulose membrane were blocked in blocking buffer (3% BSA in PBST) at 4 °C overnight. Then, 0.5 μg/mL GST-tagged proteins in blocking buffer were incubated with the membranes for 1 h at room temperature. The membranes were washed three times with PBST, and then incubated with anti-GST (1:2000 in blocking buffer) for 1 h at room temperature. Further, the membranes were washed three times with PBST, and then incubated with second antibody (1:8000 in blocking buffer) for 1 h at room temperature. Chemiluminescence were detected using GE Healthcare Amersham ECL kit.

### Immunoprecipitation-Mass Spectrometry (IP-MS) and phosphorylation analysis

The following buffers were prepared for carrying out IP-MS on LAZY4-GFP. Lysis buffer: 1x RIPA buffer (Abcam: ab156034), 1 mM PMSF, 1x protease inhibitor cocktail (Mei5bio: MF182-plus-01), and 1x phosphatase inhibitor cocktail (Mei5bio: MF183- 01). Dilution buffer: 50 mM Tris-HCl (pH 7.5), 1 mM Beta-glycerophosphate, 1 mM Sodium orthovanadate, 0.5% Sodium deoxycholate, 1mM EGTA, 1mM EDTA, and 150 mM Sodium chloride. Wash buffer: 100 mM Tris-HCl (pH 7.5), 10 mM EDTA, 10 mM EGTA, and 5 mM KCl. Four-day-old *Arabidopsis* seedlings (around 4 g) were collected and ground into powder with liquid nitrogen, and proteins were extracted using lysis buffer and diluted to around 1 mg/mL with dilution buffer. Immunoprecipitation was then performed with GFP-Trap beads (ChromoTek, gtma-20), using 50 μL GFP-Trap beads for around 6 mg proteins. Proteins bound on the beads could be collected using one of the following two strategies. First, the proteins were eluted from the beads with 50-60 μL 0.2 M glycine solution (pH 2.5) by gently shaking for 30-60 s, followed by magnetic or centrifuging separation. Then, the supernatant was transferred into a new centrifuge tube and 10% (v/v) Tris-base (pH 10.4) was added into the tube. Second, the beads were resuspended in 100 μL 2x SDS loading buffer and boiled for 10 min at 100 °C, and the proteins in the supernatant were separated by SDS-PAGE. The gel was stained with silver stain kit (Thermo: 24612), and the target bands were collected. The protein samples were analyzed on an Orbitrap Fusion Lumos Tribrid Mass Spectrometer (Thermo).

To study the phosphorylation levels of LAZY4 proteins, the data collected by tandem MS spectrometry were analyzed by SEQUEST HT (Proteome Discoverer) or Andromeda (Maxquant). The phosphorylation sites were identified based on MS2 results. MS1 peak area intensities of phosphorylated peptides were calculated for quantification, normalized by the total area intensity of phosphorylated and non-phosphorylated peptides.

### GST pull-down

Recombinant His-tagged MPK3 (∼10 μg) was activated by incubation with recombinant His-MKK5^DD^ (∼1 μg, mutated MKK5 with constitutive activity) in the presence of 50 μM ATP in 50 μL of reaction buffer (25 mM Tris, pH 7.5, 10 mM MgCl_2_ and 1mM DTT) at 37 °C for 30 min. These proteins were mixed with GST-LAZY4ΔCCL (∼100 μg) at 4 °C overnight, and then incubated with Glutathione Sepharose 4B (GE Healthcare) for 3 h at 4 °C. Alternatively, these proteins were mixed with GST-LAZY4ΔCCL and incubated with Glutathione Sepharose 4B (GE Healthcare) at 4 °C overnight. Resins were washed three times with wash buffer (First, 50 mM Tris-HCl pH 7.5, 150 mM NaCl; Second, 50 mM Tris-HCl pH 7.5, 200 mM NaCl; Third, 50 mM Tris-HCl pH 7.5, 250 mM NaCl). Pellet proteins were analyzed by Western blot using anti-His and anti-GST antibodies.

### ***In vitro* kinase assay**

Recombinant GST-tagged MPK3 (10 μg) was activated by incubation with recombinant GST-MKK5^DD^ (1 μg) in the presence of 50 μM ATP in 50 μL of reaction buffer (25 mM Tris, pH 7.5, 10 mM MgCl_2_ and 1mM DTT) at 37 °C for 30 min. Then, GST-LAZY4ΔCCL (∼100 μg) was added and reacted in the same reaction buffer with 10 mM ATP at 37 °C for 60 min. Equal volumes of 2x SDS loading buffer were added and boiled at 95 °C for 10 min to stop the reaction. Phosphorylation of LAZY proteins was analyzed by mass-spectrometry and Western blot.

### Antibody generation

We generated a polyclonal LAZY4 antibody designated anti-LAZY4. The synthetic peptide KQEITHRPSISSASSHHPR derived from the *Arabidopsis* LAZY4 was injected into rabbits as antigen, using 400 μg for the first injection and 200 μg for the following three times. Polyclonal anti-LAZY4 antibodies were purified from rabbit serum using the synthetic peptide by affinity chromatography.

### Yeast two hybrid assays

The yeast two-hybrid assays were performed as described previously^61^. The coding sequences of TOC33, TOC34, TOC64, TOC75 (TOC75-III), TOC90, TOC120, TOC132, TOC132-GM, TOC159-GM, LAZY4, LAZY4-D5/9/13 and LAZY4-A5/9/13 were amplified using the primers listed in Table S3, and the mutation sites are listed in Table S1. The TOC fragments were subcloned into the pLexA vector, and LAZY4, LAZY4-D5/9/13 and LAZY4-A5/9/13 were subcloned into the pB42AD vector. The LexA fusion constructs (bait) and the activation domain fusion constructs (prey) were co-transformed into yeast strain EGY48[p8op-lacZ] (CLONTECH). Clones containing both constructs were selected on medium lacking His, Trp and Ura, and transferred onto minimal SD/Gal/Raf/-His-Trp-Ura agar plates containing X-gal for testing β-Galactosidase activity^62^.

## Notes

### Competing Interest Statement

The authors have declared no competing interest.

