## Supplementary Figures and Tables for "Amyloplast sedimentation repolarizes LAZYs to achieve gravity sensing in plants"

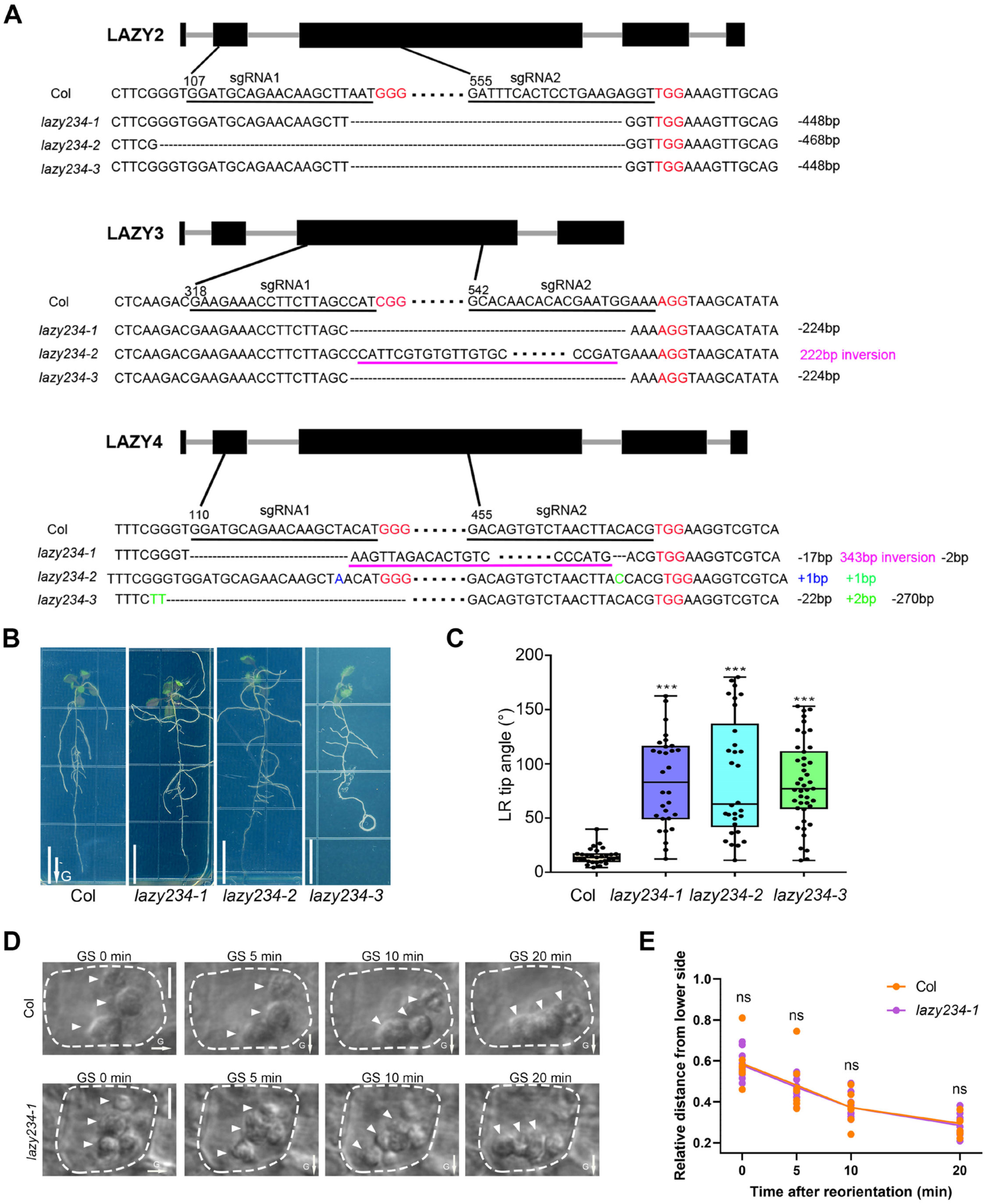


**Figure S1.** ***Arabidopsis LAZY2*****,** ***3*** **and** ***4*** **are essential for lateral root gravitropism, while they have no effect on amyloplast sedimentation, related to Figure 1**

(**A**) Gene structures and mutation sites of *Arabidopsis LAZY2*, *LAZY3* and *LAZY4* in the three alleles of *lazy2 lazy3 lazy4* (*lazy234*) triple mutants. For each LAZY gene, two sgRNAs were designed to edit the genome. PAM sequences are labeled in red, and sgRNA sequences are underlined adjacent to the PAM in Col. The mutation details are shown for each allele, including deletion, insertion and inversion.

(**B** and **C**) *Arabidopsis* lateral roots lacking *LAZY2*, *3* and *4* were agravitropic. (B) Seedlings were grown vertically on MS plates under white light, and the primary roots were aligned to the gravity vector on day 7 and continued to grow until day 16. Scale bars, 1 cm. Arrows labeled "G" indicate the direction of gravity. (C) The growth angles of lateral root (LR) tips from the gravity vector were measured on day 16. T-test was used to assess the statistical significance by comparing the three *lazy234* mutants to wild type (***$p < 0.001$; $n \geqslant 30$ for each genotype). (**D** and **E**) Amyloplast sedimentation in Col and *lazy234* is similar. (D) Four-day old seedlings were grown on MS plates under white light. Then, amyloplasts in the columella cells of wild-type and *lazy234* seedlings after 90° reorientation were imaged using a microscope with a vertical stage. Bars, 5 μm. Arrowheads indicate the positions of amyloplasts. (E) The relative distances from the amyloplasts to the new bottom of columella cells after 90° reorientation were measured at several time points. The average relative-distances of several amyloplasts within a cell was used to indicate the relative position of the amyloplasts in the cell. The distance from the top to the bottom of the cells was set as 1. The T-test was performed for the analysis. ns represents no significant difference between Col and *lazy234* mutant ($P > 0.05$; $n \geqslant 7$).

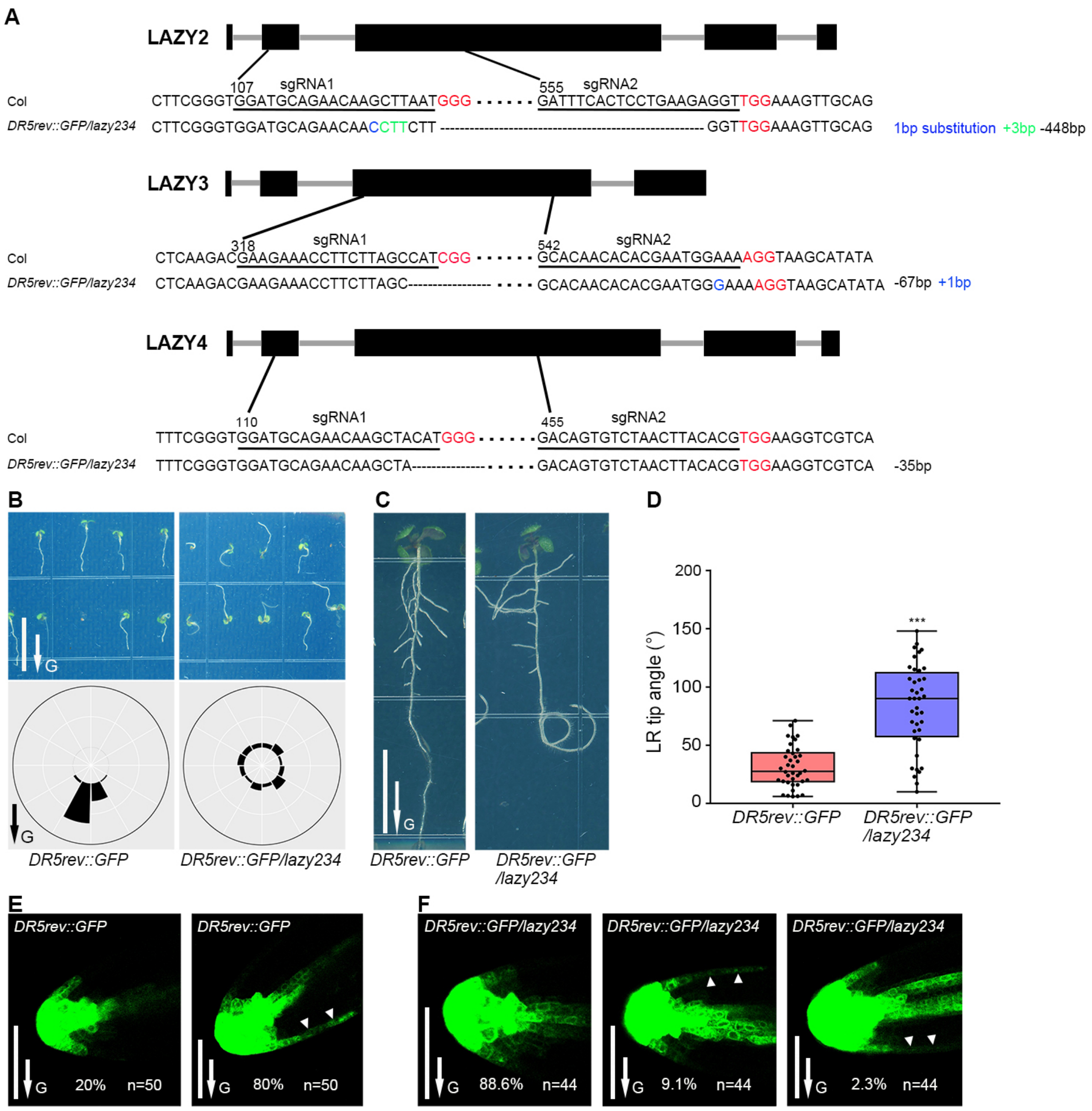


**Figure S2. Both the primary and lateral roots of *DR5rev::GFP/lazy234* lost gravitropic responses, and gravistimulation-induced asymmetrical distribution of auxin was disrupted in their lateral roots, related to Figure 1**

(**A**) Gene structures and mutation sites of *Arabidopsis LAZY2, LAZY3* and *LAZY4* in the *DR5rev::GFP/lazy234*. For each *LAZY* gene, two sgRNAs were designed to edit the genome. PAM sequences are labeled in red, and sgRNA sequences are underlined adjacent to the PAM in Col. The mutation details are shown for each allele, including deletion, insertion and substitution.

(**B**) Growth orientation of *DR5rev::GFP/lazy234* primary roots. Seedlings were grown vertically under white light for 4 days. Frequencies of root angles in each 30° division around a circle are shown (n>50 for each genotype).

**(C** and **D)** Growth orientation of *DR5rev::GFP/lazy234* lateral roots. Seedlings were grown vertically under white light, and the primary roots were aligned to the gravity vector on day 7.

The growth angles of lateral root (LR) tips from the gravity vector were measured on day 13. T-test was used to assess the statistical significance by comparing *DR5rev::GFP/lazy234* to *DR5rev::GFP* (***p < 0.001; n>35 for each genotype).

**(E)** Expression of *DR5rev::GFP* in the stage 2 lateral roots of wild type. Two categories of GFP fluorescence patterns were observed and the proportions of each category are indicated. Left, symmetric distribution; Right, higher accumulation on the lower side. Arrowheads indicate the asymmetric accumulation of auxin.

**(F)** Expression of *DR5rev::GFP* in the stage 2 lateral roots of *lazy234*. Three categories of GFP fluorescence patterns were observed and the proportions of each category are indicated. Left, symmetric distribution; Middle, higher accumulation on the upper side. Right, higher accumulation on the lower side. Arrowheads indicate the asymmetric accumulation of auxin.

In B, C, E and F, arrows labeled “G” indicate the direction of gravity. In B and C, scale bars, 1 cm; in E and F, scale bars, 50 μm.

54

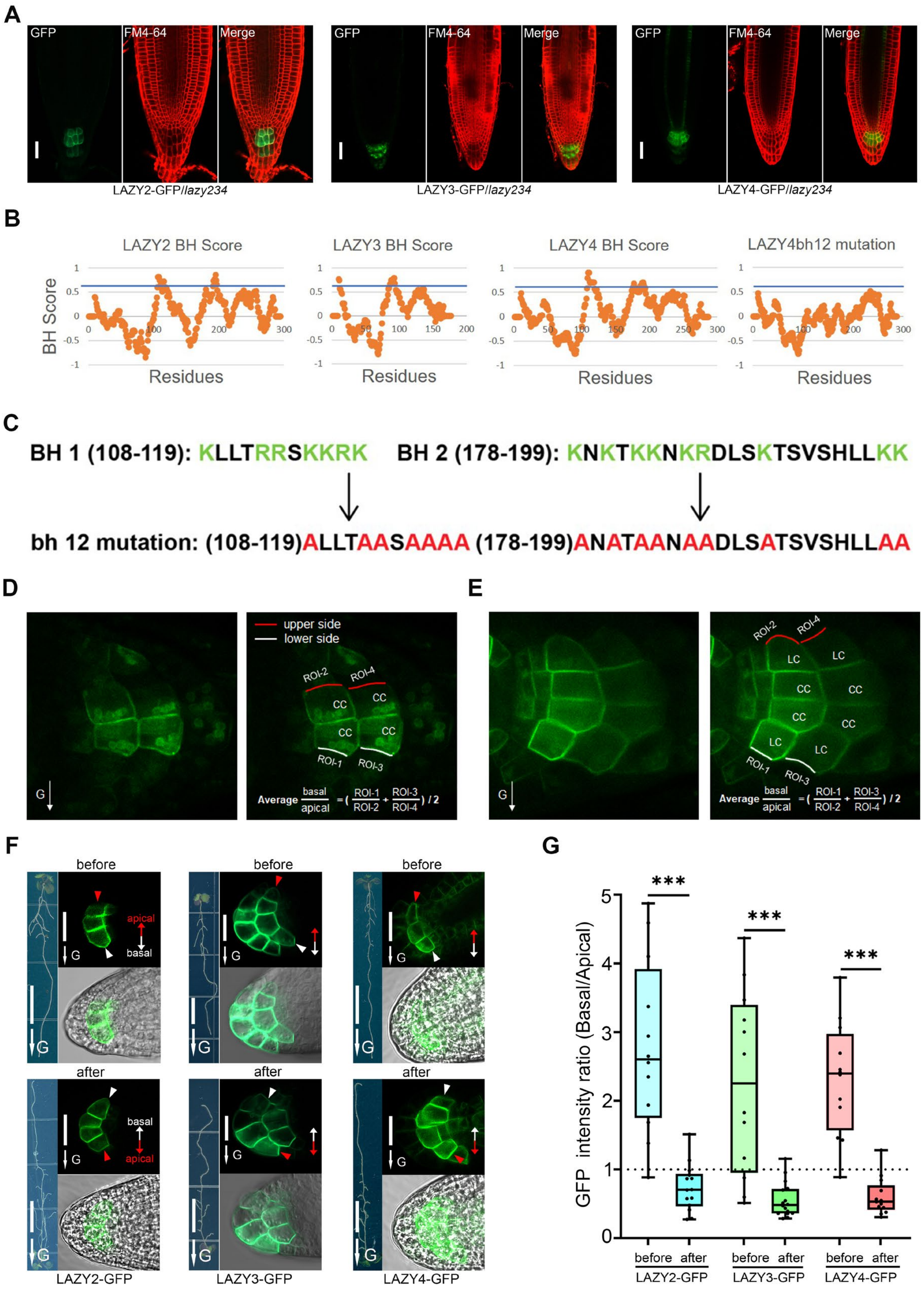
A
GFP
FM4-64
Merge
LAZY2-GFP/lazy234
LAZY3-GFP/lazy234
LAZY4-GFP/lazy234
B
LAZY2 BH Score
LAZY3 BH Score
LAZY4 BH Score
LAZY4bh12 mutation
BH Score
Residues
C
BH 1 (108-119): KLLTRRSKKRK
BH 2 (178-199): KNKTKKNKRDLSKTSVSHLLKK
bh 12 mutation: (108-119)ALLTAASAAAA (178-199)ANATAANAADLSATSVSHLLAA
D
E
upper side
lower side
ROI-1
ROI-2
ROI-3
ROI-4
CC
LC
G
Average basal/apical = (ROI-1/ROI-2 + ROI-3/ROI-4) / 2
F
before
after
apical
basal
LAZY2-GFP
LAZY3-GFP
LAZY4-GFP
G
GFP intensity ratio (Basal/Apical)
***
before after
LAZY2-GFP
LAZY3-GFP
LAZY4-GFP

55

**Figure S3. The localization of LAZY2, 3 and 4 proteins in the columella cells of *Arabidopsis* primary and lateral roots, related to Figures 2 and 3**

(**A**) The localization of *Arabidopsis* LAZY2, LAZY3 and LAZY4 proteins in root tips. Four-day old seedlings were grown on MS plates under white light, and then imaged by confocal microscope. The root tips were treated with 5 μM FM4-64 for 2 minutes, and the fluorescence of LAZY-GFP proteins and FM4-64 were collected from the root tips. Scale bars, 20 μm. The merged images are the same as those in the Figures 2A to 2C, whereas the images of separated channels are shown here for better view of the GFP signals.

(**B**) BH score predictions of LAZY2, LAZY3, LAZY4 and LAZY4 with bh sites mutation (LAZY4bh12). Predictions were made on https://hpcwebapps.cit.nih.gov/bhsearch/. Blue grid line refers to 0.6, which is a threshold for possible PtdIns binding.

(**C**) The schematic of LAZY4bh12 mutating K/R to A within two basic hydrophobic peaks.

(**D** and **E**) Measurement of LAZY-GFP fluorescence intensity ratio. Images showing two representative patterns of LAZY-GFP fluorescence in columella cells, with clear signals only in central columella (CC) cells (**B**), or in both of central columella (CC) cells and lateral columella (LC) cells (**C**). In each panel, the left is the original image, and the right image shows how the intensity ratio between basal and apical is calculated. Upper or lower side of the plasma membrane in columella cells were selected as region of interest (ROI) for measurement of LAZY-GFP fluorescence intensity. Signal ratio (basal/apical) for one root was calculated as an average of signal intensity ratios within this root.

(**F** and **G**) LAZY2, 3 and 4 proteins show polar distribution in the columella cells of *Arabidopsis* lateral roots under the regulation of gravity. (D) LAZY2, LAZY3 and LAZY4 proteins are mainly accumulated on the lower side of columella cells in the lateral roots (LRs) at the stage 2 under both regular growth (upper) or after 180° reorientation (lower). The primary roots of *ProLAZY2:LAZY2-GFP* (LAZY2-GFP/Col), *ProLAZY3:LAZY3-GFP/lazy234* (LAZY3-GFP) and *ProLAZY4:LAZY4-GFP/lazy234* (LAZY4-GFP) growing under white light conditions were aligned to the gravity vector on the $7^{th}$ day and continued to grow until the $12^{th}$ day. The fluorescence of LAZY-GFP was collected before (Upper in each panel), or after the seedlings were reoriented 180° and grown for another 6 h (Lower in each panel). Double-headed arrows show the apical-basal axis of the plant bodies. Red and white arrow heads indicate the accumulation of LAZY-GFP proteins on the apical and basal sides of columella cells respectively. For the whole seedlings, scale bars, 1 cm; for all the others, scale bars, 20 μm. Arrows labeled “G” indicate the direction of gravity. (E) The statistical analysis of LAZY-GFP fluorescence intensity ratios of the two sides of the lateral roots before and after $180^{o}$ rotation. Asterisks indicate Student’s t-test values (***, $P < 0.001$; $n \geqslant 12$).

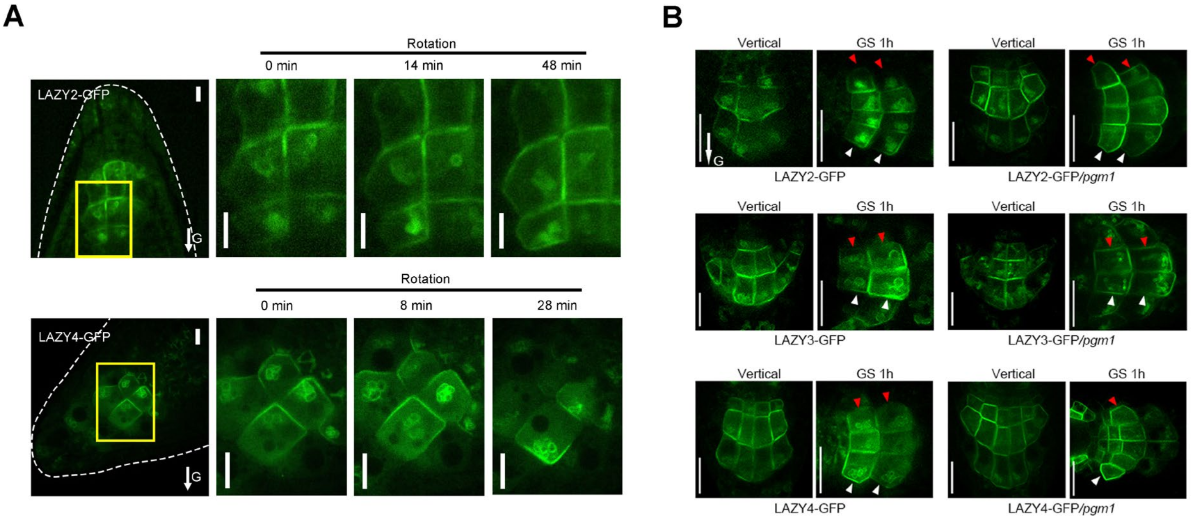


**Figure S4. Amyloplast sedimentation promotes the redistribution of LAZY proteins, related to Figure 3**

**(A)** Gravity-triggered redistribution of LAZY2 and LAZY4 proteins to the lower side of columella cells correlates with the sedimentation of amyloplasts. The seedlings of LAZY2-GFP (without CHX treatment) and LAZY4-GFP (with 10 µM CHX treatment to inhibit new protein synthesis) were reoriented to a position with amyloplasts on the top of the columella cells, and kept for fluorescence collection at several time points. The area inside yellow box was zoomed to show details of fluorescence. White dash lines indicate the contour of root tips. Scale bars, 10 µm.

**(B)** Mutation of *PGM1* delayed the redistribution of LAZY2-GFP, LAZY3-GFP and LAZY4-GFP proteins in root columella cells. Seedlings were grown vertically and then reoriented 90° and kept 1 h for gravistimulation (GS 1h). Red and white arrowheads indicate the accumulation of LAZY-GFP proteins on the upper and lower sides of columella cells respectively. Scale bars, 20 µm. Statistical data are shown in Figures 3M to 3O.

In A and B, arrows labeled "G" indicate the direction of gravity.

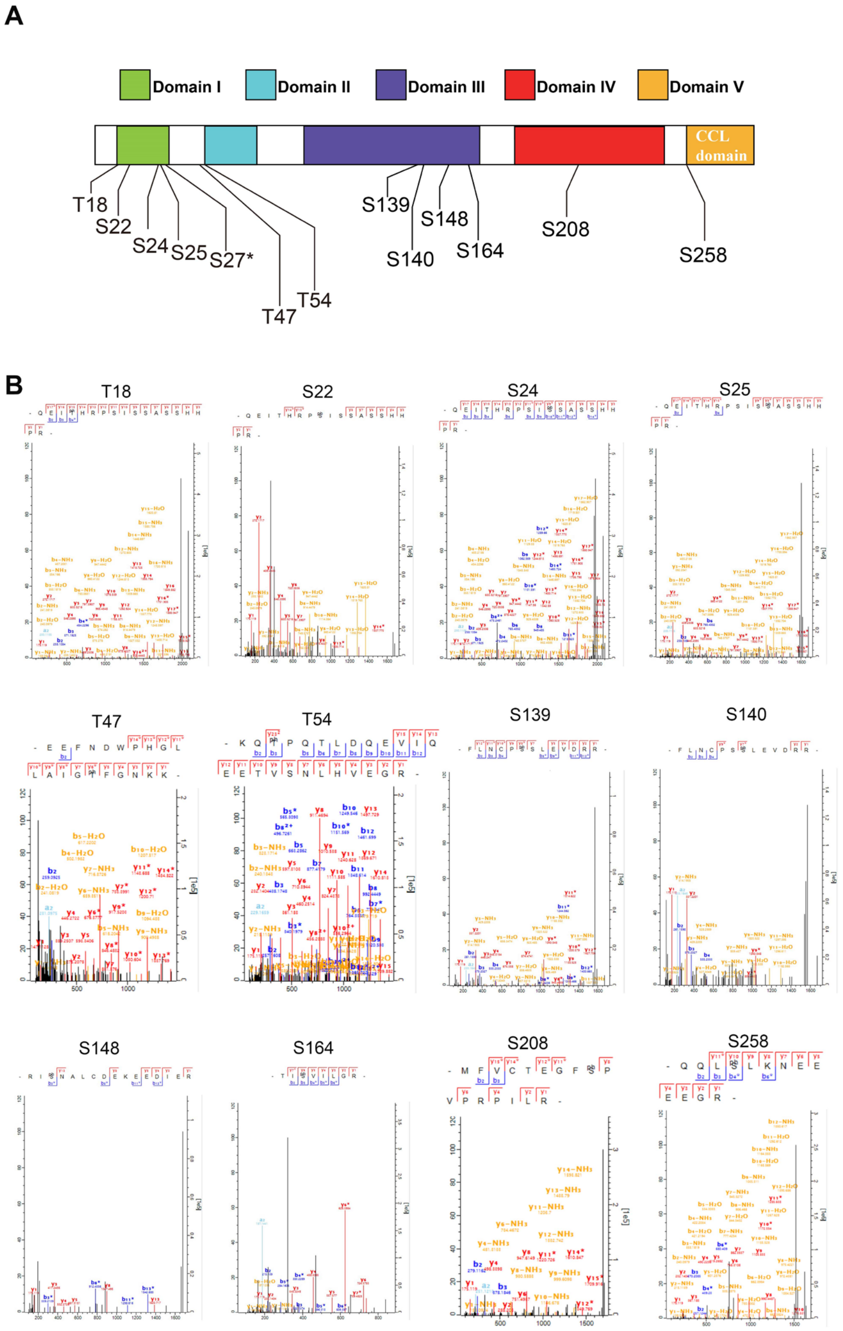


**Figure S5. Phosphorylation sites of LAZY4 protein under gravistimulation, related to Figure 4**

(**A**) Four-day-old LAZY4-GFP seedlings were grown vertically or re-orientated 90$^o$ (gravistimulation), and LAZY4-GFP proteins were immunoprecipitated for Mass-Spectrometric (MS) analysis. Phosphorylation sites identified by MS are shown. The star “*” used to label S27 means that S27 is possibly phosphorylated but not fully confirmed based on the MS/MS spectra.

(**B**) MS/MS spectra. X axis shows the m/z value of each fragment ion. Y axis shows the relative abundance (left) or ion intensity (right) of the corresponding daughter ions. Peptide sequences are displayed above the spectra.

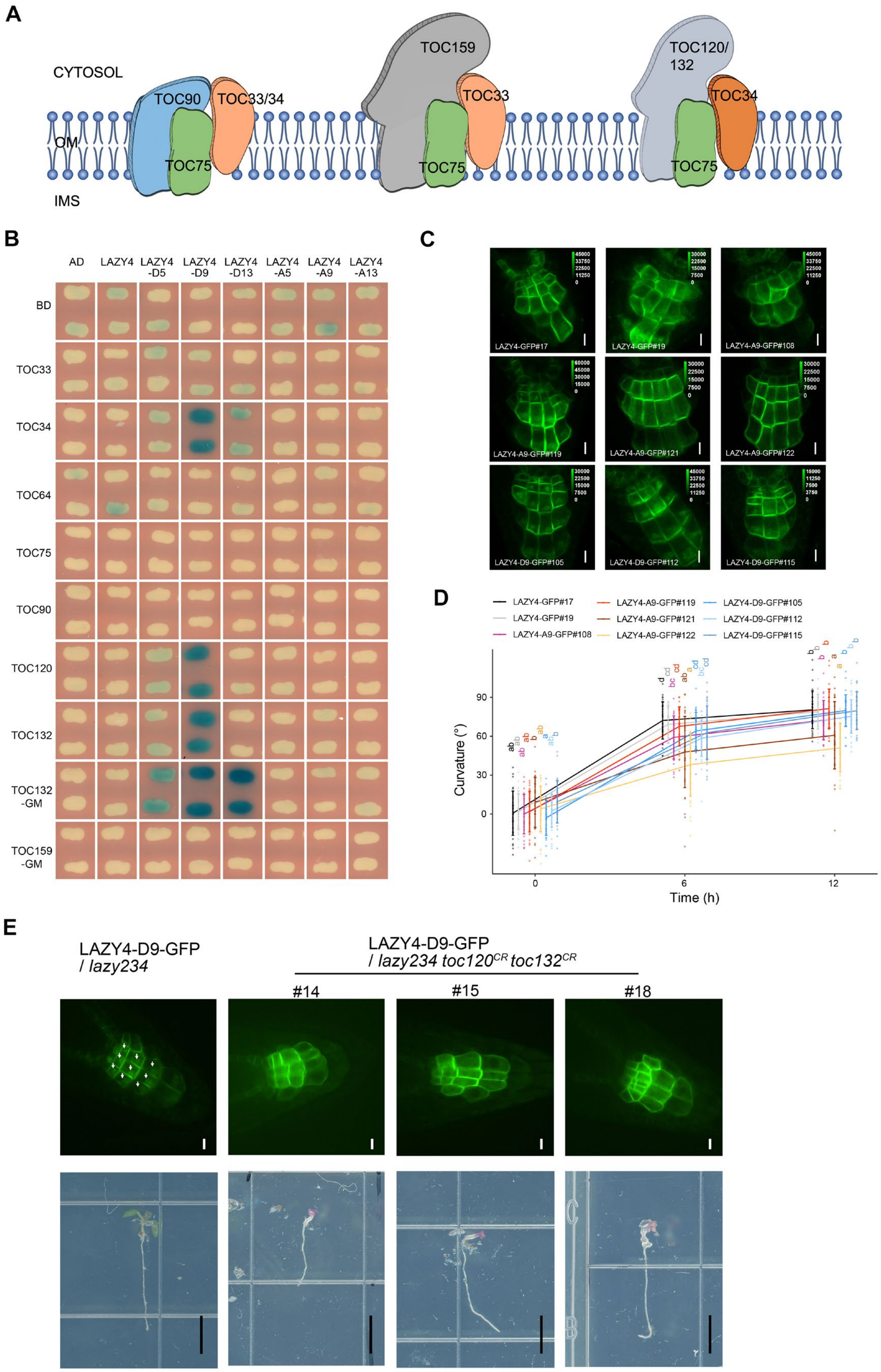
A
CYTOSOL
OM
IMS
TOC90
TOC33/34
TOC75
TOC159
TOC33
TOC75
TOC120/
132
TOC34
TOC75
B
AD
LAZY4
LAZY4
-D5
LAZY4
-D9
LAZY4
-D13
LAZY4
-A5
LAZY4
-A9
LAZY4
-A13
BD
TOC33
TOC34
TOC64
TOC75
TOC90
TOC120
TOC132
TOC132
-GM
TOC159
-GM
C
LAZY4-GFP#17
LAZY4-GFP#19
LAZY4-A9-GFP#108
LAZY4-A9-GFP#119
LAZY4-A9-GFP#121
LAZY4-A9-GFP#122
LAZY4-D9-GFP#105
LAZY4-D9-GFP#112
LAZY4-D9-GFP#115
D
LAZY4-GFP#17
LAZY4-GFP#19
LAZY4-A9-GFP#108
LAZY4-A9-GFP#119
LAZY4-A9-GFP#121
LAZY4-A9-GFP#122
LAZY4-D9-GFP#105
LAZY4-D9-GFP#112
LAZY4-D9-GFP#115
Curvature (°)
90
60
30
0
0
6
12
Time (h)
E
LAZY4-D9-GFP
/ lazy234
LAZY4-D9-GFP
/ lazy234 toc120CR toc132CR
#14
#15
#18

**Figure S6. Effects of phosphomimic or phosphodead LAZY4 proteins, related to Figure 5**

**(A)** Distinct TOC complexes in *Arabidopsis*. In *Arabidopsis* TOC complexes, TOC75 is a central pore component localized within the outer membranes of plastids, and other TOC proteins are receptors facing the cytosol. TOC33 and TOC159 are components of a class of TOC complexes that specifically import photosynthesis-related preproteins. TOC34, TOC120 and TOC132 are components of another class of TOC complexes that specifically import non-photosynthetic or house-keeping preproteins. TOC90 may be partially redundant with TOC159, and associates with TOC33 or TOC34. Abbreviations: TOC, Translocons at the Outer envelope membrane of Chloroplasts; OM, Outer Membrane; IMS, Intermembrane Space. The models of TOC complexes were drawn based on information from a previous review[37].

(**B**) Yeast two-hybrid assays to test interactions between LAZY4 variants and TOC proteins. Interactions between LAZY4 variants and TOC proteins were examined by yeast growth on media with X-Gal but lacking Trp, His and Ura. The phosphorylation sites of LAZY4 were mutated to Asp (D) or Ala (A) as indicated in Table S1. Blue indicates positive interactions. Reporter: LacZ; substrate, X-Gal. These results were from the same experiment shown in Figure 5A, but with longer growth time (40 h) and more LAZY4 variants (mutated to A).

**(C)** Fluorescence in the root tip of LAZY4-GFP/*lazy234,* LAZY4-A9-GFP/*lazy234* and LAZY4-D9-GFP/*lazy234*. The seedlings were grown vertically for four days. Fluorescence of LAZY4-GFP, LAZY4-A9-GFP and LAZY4-D9-GFP in columella cells was collected by confocal microscopy. All images were acquired by excitation light (488 nm laser) of same intensity except for LAZY4-A9-GFP#121 (lower laser intensity), LAZY4-A9-GFP#122 (lower laser intensity), LAZY4-D9-GFP#112 (higher laser intensity). Scale bars, 10 μm.

(**D**) Statistical analysis of LAZY4-GFP/*lazy234,* LAZY4-A9-GFP/*lazy234* and LAZY4-D9-GFP/*lazy234* phenotypes. Seedlings were grown vertically for three days under regular white light (70 $\mu M/m^2/s$) and reoriented 90° for gravistimulation under weak light (15 $\mu M/m^2/s$). At each time point, root curvatures of the transgenic lines were compared. One way ANOVA was used to assess the statistical significance, and the homogeneous subsets were calculated by Waller-Duncan multiple comparison test.

(**E**) Simultaneous mutation of TOC120 and TOC132 disrupts the localization of LAZY4-D9-GFP onto the surface of amyloplasts. The LAZY4-D9-GFP/*lazy234* $toc120^{CR}$ $toc132^{CR}$ seedlings were selected based on the color of cotyledons (chlorophyll deficiency) from T2 populations using CRISPR to mutate TOC 120 and TOC132 in LAZY4-D9-GFP/*lazy234*. White arrows indicate localization of LAZY4-D9-GFP on the amyloplasts. Upper, scale bars, 10 μm; Lower, scale bars, 0.3 cm.

**Table S1. Phosphorylation of LAZY4 proteins under gravistimulation or in *in vitro* kinase assay**

| Peptides | MS1 intensity of phosphorylated peptide | up-regulated upon gravistimulation | Phosphorylation sites | IP-MS | | | | | | | in vitro kinase assay (MPK3) | | Included in | | |
|---|---|---|---|---|---|---|---|---|---|---|---|---|---|---|---|
| | | | | PR1 | PR2 | PR3 | PR4 | R1 | R2 | R3 | KR1 | KR2 | A5/D5 | A9/D9 | A13/D13 |
| QEITHRPSISSASSHHPR | +++ | √ | T18 | | | √ | √ | √ | √ | √ | √ | √ | √ | √ | √ |
| | | | S22 | | | * | √ | √ | √ | √ | | | √ | √ | √ |
| | | | S24 | | | * | √ | √ | √ | √ | | √ | √ | √ | √ |
| | | | S25 | | | * | √ | √ | √ | √ | | | √ | √ | √ |
| | | | S27 | | | | * | * | * | | | | √ | √ | √ |
| EEFNDWPHGLLAIGTFGNK/ EEFNDWPHGLLAIGTFGNKK | ++ | √ | T47 | | | | | √ | √ | √ | √ | √ | | | √ |
| KQTPQTLDQEVIQEETVSNL HVEGR | + | √ | T54 | | | | | √ | √ | √ | √ | √ | | | √ |
| FLNCPSSLEVDR/ FLNCPSSLEVDRR | ++ | √ | S139 | √ | √ | | | √ | √ | √ | | √ | | √ | √ |
| | | | S140 | √ | √ | | | √ | √ | √ | | √ | | √ | √ |
| RISNALCDEKEEDIER | + | | S148 | √ | √ | √ | | | √ | √ | √ | | | √ | √ |
| TISVILGR | + | | S164 | √ | | | | √ | √ | √ | | | | √ | √ |
| MFVCTEGFSPVPRPILR | ++ | √ | S208 | | | | | √ | √ | √ | √ | √ | | | √ |
| QQLSLKNEEEEGR | + | √ | S258 | | | | | √ | | | | √ | | | √ |

This table summarizes the mass spectrometry results of the IP-MS experiment shown in Figure 4A (three replicates: R1, R2, and R3) and four preliminary replicates (PR1 to PR4), and in vitro kinase assays shown in Figure 4E (two replicates: KR1 and KR2). For the MS1 intensity of phosphorylated peptides based on R1, R2 and R3: +++, above $1\times10^8$; ++, between $1\times10^7$ and $1\times10^8$; +, below $1\times10^7$. If max MS1 intensity of a phosphorylated peptide in samples of gravistimulation was 5-fold greater than the control under vertical condition in at least two experimental replicates, the phosphorylation of the peptide is recorded as "up-regulated upon gravistimulation". The phosphorylation sites with explicit MS/MS spectra are labeled " √ ", and possible phosphorylation sites are labeled "*". For LAZY4-A5/D5, A9/D9, and A13/D13 mutagenesis, selected amino acids were mutated to Ala (A) or Asp (D). For A5/D5, the 5 phosphorylated amino acids identified in a phosphorylated peptide with the highest MS1 intensity. For A9/D9, the total 9 phosphorylated amino acids identified in PR1 to PR4, which were also identified in R1, R2 and R3. For A13/D13, all the 13 phosphorylated amino acids identified in R1, R2 and R3.

**Table S2. Mutation sites of *Arabidopsis TOC120* and *TOC132* in the LAZY4-GFP/*lazy234 toc120*$^{CR}$ *toc132*$^{CR}$ seedlings**

| genotype | *TOC120* | *TOC132* |
|---|---|---|
| Type 1 | Insert 13 bp (1753), delete 7 bp (1753-1759), delete 6 bp (2727-2732). | Delete I1245 bp (1694-2938), insert 5 bp (1964), A2943C. |
| Type 2 | Delete 15bp (1744-1758). | Delete 1245 bp (1694-2938), insert 5 bp (1964), A2943C. |
| Type 3 | Delete 84 bp (2226-2309), the sequencing result was bimodal. | Delete 1245 bp (1694-2938); insert 5 bp (1964); A2943C. |
| Type 4 | Insert 56 bp(2105), delete 38 bp (2105-2142), delete 39 bp (2157-2195), the sequencing result was bimodal. | Delete 1245 bp (1694-2938); insert 5 bp (1964); A2943C. |
| Type 5 | Insert 13 bp (1753), delete 7 bp (1753-1759), delete 6 bp (2727-2732). | Delete 1180 bp (1694-2873). |
| Type 6 | Delete 135 bp (2227-2361), delete 28 bp (2388-2415), the sequencing result was bimodal. | Delete 1180 bp (1694-2873). |
| Type 7 | The sequencing result was bimodal. | Delete 1180 bp (1694-2873). |
| Type 8 | Insert 13 bp (1753), delete 7 bp (1753-1759), delete 6 bp (2727-2732). | Delete 1179 bp (1696-2874). |
| Type 9 | Delete 27 bp (2043-2069), delete 42 bp (2105-2146), the sequencing result was bimodal. | Delete 1179 bp (1696-2874). |
| Type 10 | Delete 88 bp (2226-2313), the sequencing result was bimodal. | Delete 1179 bp (1696-2874). |
| Type 11 | The sequencing result was bimodal. | Delete 1179 bp (1696-2874). |
| Type 12 | Delete 15bp (1744-1758). | Delete 1178 bp (1696-2873). |
| Type 13 | Insert 13 bp (1753), delete 7 bp (1753-1759), insert 18 bp (2369), delete 11 bp (2369-2379)，the sequencing result was bimodal. | Delete 1178 bp (1696-2873). |

**Table S3. Primers used in this study**

| Purpose | Primer name | Sequence (5'-3') |
|---|---|---|
| LAZY-sgRNA constructs | LAZY2-sgRNA1-F | GATTGGATGCAGAACAAGCTTAAT |
| | LAZY2-sgRNA1-R | AAACATTAAGCTTGTTCTGCATCC |
| | LAZY2-sgRNA2-F | GATTGATTTCACTCCTGAAGAGGT |
| | LAZY2-sgRNA2-R | AAACACCTCTTCAGGAGTGAAATC |
| | LAZY3-sgRNA1-F | GATTGAAGAAACCTTCTTAGCCAT |
| | LAZY3-sgRNA1-R | AAACATGGCTAAGAAGGTTTCTTC |
| | LAZY3-sgRNA2-F | GATTGCACAACACACGAATGGAAA |
| | LAZY3-sgRNA2-R | AAACTTTCCATTCGTGTGTTGTGC |
| | LAZY4-sgRNA1-F | GATTGGATGCAGAACAAGCTACAT |
| | LAZY4-sgRNA1-R | AAACATGTAGCTTGTTCTGCATCC |
| | LAZY4-sgRNA2-F | GATTGACAGTGTCTAACTTACACG |
| | LAZY4-sgRNA2-R | AAACCGTGTAAGTTAGACACTGTC |
| Genotype the mutants generated by CRISPR | LAZY2-genotyping-F | GATTCATCAATAACTCGAGTT |
| | LAZY2-genotyping-R | AAGAGATTTTATCGTTGTGTC |
| | LAZY3-genotyping-F | CTTGGAGAACATTGACAGAAA |
| | LAZY3-genotyping-R1 | ATTCACTTGGTTAATTACACC |
| | LAZY3-genotyping-R2 | CCATGACTATCAATCAGGTCCA |
| | LAZY4-genotyping-F | CACCTTAAACCCGAGTTATTT |
| | LAZY4-genotyping-R | AAAGATTCACTGATCAGGTCA |
| | TOC132-sgGM-GT-F | GCAAACCCCGCATAATGTTG |
| | TOC132-sgGM-GT-R | GTCACTTGACCTGAGACAGA |
| | TOC120-sgGM-GT-F | TCCGCCTGACATTGTGTTAT |
| | TOC120-sgGM-GT-R | ACTGAATCACCTAAGAGCGT |
| | Cas9-test-F | AAGATCGAGAAGATCCTGACCTT |
| | Cas9-test-R | CATAGGTTTTCAGCCGTTCCT |
| *ProLAZY:LAZY-GFP* constructs | ProLAZY2-F | CATGCCTGCAGGATTGCGTGACGATTGAGCACC |
| | LAZY2-R | CCTCTAGACGCCGCCGCCGCCGCTATCTCGAGAACTATGAC |
| | ProLAZY3-F | GCATGCCTGCAGGTTAGTTAAATTATAAAAAAAAATCGGTTTTG |
| | LAZY3-R | CCGCTCGAGCGCCGCCGCCGCCGCCATCTCCAAGACAATATCTG |
| | ProLAZY4-F | ATGCCTGCAGGCGATTAGCGAATGTTTAAGGCAACG |
| | LAZY4-R | GACTAGTCGCCGCCGCCGCCGCGATCTCAAGAACAATGACTG |
| Genotype *pgm1* | pgm1-F | GTCATTTAAAGTTTCTTGGATGATTTACAATGC |
| | pgm1-R | CTTACCTCCATGAGCTCCAAATAGTC |
| *ProLAZY4:LAZY4-D9-GFP* constructs | Gbs-LAZY4pro-D9-F | GCTTGCATGCCTGCAGGCGATTAGCGAATGTTTAAGG |
| | LAZY4-D9-mut-F1 | ATGAAGTTTTTCGGGTGG |
| | LAZY4-D9-mut-R1 | TCAGCATCATCTATATCTGGTCTATGAT |

| | | |
|---|---|---|
| | | CAATCTCTTGTTTCCCATG |
| | LAZY4-D9-mut-F2 | GATCATAGACCAGATATAGATGATGCT GATTCTCATCATCCGAGAGA |
| | LAZY4-D9-mut-R2 | TCGATTGTACGCTCAATGTCTTCCTCCTT CTCATCACAAAGCGCGTTATCGATTCTT CTATCGACCTCAAGATCATCAGGACAAT TCAAGAATCTA |
| | LAZY4-D9-mut-F3 | GATGATCTTGAGGTCGATAGAAGAATC GATAACGCGCTTTGTGATGAGAAGGAG GAAGACATTGAGCGTACAATCGATGTT ATCCTAGGGAGATGC |
| | LAZY4-D9-mut-R3 | CGCCGCCGCCGCCGCCGC |
| | Gbs-LAZY4pro-D9-R | AAAAACTTCATGTCTTAGTGACCCGGAA G |
| | Gbs-LAZY4-D9-F | GTCACTAAGACATGAAGTTTTTCGGGTG |
| | Gbs-LAZY4-D9-R | GCCCTTGCTCACCATACTAGTCGCCGCC GCCGC |
| | pJIM-sequence-F | CTGATAGTTTAAACTGAAGGCGGG |
| | pJIM-sequence-R | GCGGCAGCGTGAAGCTG |
| *ProLAZY4:LAZY4-A9-GFP* constructs | Gbs-LAZY4pro-A9-F | TGAATTAAGCTTGCATGCCTGCAGGCGA TTAGCGAATGTTTAAGG |
| | Gbs-LAZY4pro-A9-R | CCGAAAAACTTCATGTCTTAGTGACCCG GAAG |
| | Gbs-LAZY4-A9-F | GTCACTAAGACATGAAGTTTTTCGGGTG |
| | Gbs-LAZY4-A9-R | TCCTCGCCCTTGCTCACCATACTAGTCG CCGCCGCCGC |
| | LAZY4-A9-mut-F1 | ATGAAGTTTTTCGGGTGG |
| | LAZY4-A9-mut-R1 | CGCAGCCGCCGCTATCGCTGGTCTATGC GCAATCTCTTGTTTCCCAT |
| | LAZY4-A9-mut-F2 | CGCATAGACCAGCGATAGCGGCGGCTG CGTCTCATCATCCGAGAGAG |
| | LAZY4-A9-mut-R2 | CGCGATTGTACGCTCAATGTCTTCCTCC TTCTCATCACAAAGCGCGTTCGCGATTC TTCTATCGACCTCAAGCGCCGCAGGACA ATTCAAGAATCT |
| | LAZY4-A9-mut-F3 | CGGCGCTTGAGGTCGATAGAAGAATCG CGAACGCGCTTTGTGATGAGAAGGAGG AAGACATTGAGCGTACAATCGCGGTTAT CCTAGGGAGATGCA |
| | LAZY4-A9-mut-R3 | CGCCGCCGCCGCCGCCGC |
| *ProLAZY3:LAZY3(truncated)-GFP* constructs | LAZY3-△N2-27-F | TTCTAGTATGAGTCCTCCTT |
| | LAZY3-△N2-27-R | AAGGAGGACTCATACTAGAA |
| | LAZY3-△CCL-F | TTTTCCTGCAGGTGTTTGCTGTTCTGGA GACG |
| | LAZY3-△CCL-R | TTTTCTCGAGCGCCGCCGCCGCCGCCGC GTTGGCGTCAACACTTGAAC |

| | | |
|---|---|---|
| Y2H constructs_LAZY4 | Gbs-LAZY4-pB42-F | CAGATTATGCCTCTCCCGAATTCATGAAGTTTTTCGGGTGGATGCAG |
| | Gbs-LAZY4-pB42-R | CGAAGAAGTCCAAAGCTTCTCGAGTCAGATCTCAAGAACAATGAAATCAGAATCTGTTTTGAC |
| | PB42AD-LAZY4-F | ATTATGCCTCTCCCGAATTCATGAAGTTTTTCGGGTGG |
| | LAZY4-A5/D5-mut-R | CAAGAATCTATCCAAAGGAAGATTC |
| | LAZY4-A5/D5-mut-F | GAATCTTCCTTTGGATAGATTCTTG |
| | LAZY4-D13-mut-R1 | ATCCTGCTTTTTGTTACCGAAATCACCAATCGCGAGTAAT |
| | LAZY4-D13-mut-F1 | GATTTCGGTAACAAAAAGCAGGATCCACAAACACTTGATC |
| | LAZY4-D13-mut-R2 | GGGATCAAAACCTTCTGTACAG |
| | LAZY4-D13-mut-F2 | TGTACAGAAGGTTTTGATCCCGTTCCTCGCCCTAT |
| | LAZY4-D13-mut-R3 | AAATCGAGCTGTTGCTTGTCT |
| | LAZY4-D13-mut-F3 | GACAAGCAACAGCTCGATTTGAAGAACGAGGAAGA |
| | LAZY4-A13-mut-R1 | CGCCTGCTTTTTGTTACCGAACGCACCAATCGCGAGTAAT |
| | LAZY4-A13-mut-F1 | GCGTTCGGTAACAAAAAGCAGGCGCCACAAACACTTGATC |
| | LAZY4-A13-mut-F2 | GTACAGAAGGTTTTGCGCCCGTTCCTCG |
| | LAZY4-A13-mut-R2 | CGAGGAACGGGCGCAAAACCTTCTGTAC |
| | LAZY4-A13-mut-R3 | CTTCAACGCGAGCTGTTGCTTGTCTT |
| | LAZY4-A13-mut-F3 | CAAGCAACAGCTCGCGTTGAAGAACGAGGAAG |
| | PB42AD-LAZY4-R | GAAGTCCAAAGCTTCTCGAGTCAGATCTCAAGAACAATGAAATCAG |
| Y2H constructs_TOC | Gbs-TOC33-plexA-F | ACGGCGACTGGCTGGAATTCATGGGGTCTCTCGTTCGTG |
| | Gbs-TOC33-plexA-R | TGGCTGCAGGTCGACTCGAGTTAAAGTGGCTTTCCACTTGTCTTG |
| | Gbs-TOC34-plexA-F | ACGGCGACTGGCTGGAATTCATGGCAGCTTTGCAAACG |
| | Gbs-TOC34-plexA-R | TGGCTGCAGGTCGACTCGAGTCAAGACCTTCGACTTGCTA |
| | Gbs-TOC64-plexA-F | ACGGCGACTGGCTGGAATTCATGGCGACCAATAATGATTTTGG |
| | Gbs-TOC64-plexA-R | TGGCTGCAGGTCGACTCGAGTCAAATAAATGCAGCAAGGGAATCC |
| | Gbs-TOC75-plexA-F | ACGGCGACTGGCTGGAATTCATGGCCGCCTTCTCCGTC |

| | | |
|---|---|---|
| | Gbs-TOC75-plexA-R | TGGCTGCAGGTCGACTCGAGTCATTGTCCATATTGCGTTTGCG |
| | Gbs-TOC90-plexA-F | ACGGCGACTGGCTGGAATTCATGAAAGGCTTCAAAGACTGGG |
| | Gbs-TOC90-plexA-R | TGGCTGCAGGTCGACTCGAGTTAGGAAACGAGAAAATTCACAATCTTTTCTTC |
| | Gbs-TOC120-plexA-F | ACGGCGACTGGCTGGAATTCATGGGAGATGGGGCTGAGATTG |
| | Gbs-TOC120-plexA-R | TGGCTGCAGGTCGACTCGAGTCAGTGTCCATATTGCATTTGCTCAGG |
| | Gbs-TOC132-plexA-F | ACGGCGACTGGCTGGAATTCATGGGAGATGGGACTGAGTTTG |
| | Gbs-TOC132-plexA-R | TGGCTGCAGGTCGACTCGAGTCATTGTCCATATTGCGTTTGCG |
| | BD-TOC132G-F | ACGGCGACTGGCTGGAATTCGGTCGTGCTTCTCCTCTTTT |
| | BD-TOC132M-R | TGGCTGCAGGTCGACTCGAGTCATTGTCCATATTGCGTTT |
| | BD-TOC159G-F | ACGGCGACTGGCTGGAATTCTCCCTAAACATACTGGTCCT |
| | BD-TOC159M-R | TGGCTGCAGGTCGACTCGAGTTAGTACATGCTGTACTTGT |
| TOC120/132-sgRNA constructs | TOC120-sgG-F | GATTGCCATCTGGTGGAGCCGAAG |
| | TOC120-sgG-R | AAACCTTCGGCTCCACCAGATGGC |
| | TOC120-sgM-F | GATTGAAGGTAGATCAACTTCCCT |
| | TOC120-sgM-R | AAACAGGGAAGTTGATCTACCTTC |
| | TOC132-sgG-F | GATTGCAGAACAGCTTGAGGCTGC |
| | TOC132-sgG-R | AAACGCAGCCTCAAGCTGTTCTGC |
| | TOC132-sgM-F | GATTGCCACTGATTGGAGGAATCA |
| | TOC132-sgM-R | AAACTGATTCCTCCAATCAGTGGC |
| *ProTOC132:TOC132-RFP* constructs | pCAMBIA1300-Sal I-TOC132g-F | CCAAGCTTGCATGCCTGCAGGTCGACTAGCTGCACCAGCTTATTGA |
| | TOC132g-R-BamH I-mRFP | TCCTCGGAGGAGGCGCTCATGGATCCTTGTCCATATTGCGTTTGCG |
| Prokaryotic expression constructs | Gbs-LAZY2-pGEX4T-1-F | CGCGTGGATCCCCGGAATTCATGAAGTTCTTCGGGTGGATGCA |
| | Gbs-LAZY2-pGEX4T-1-R | TCACGATGCGGCCGCTCGAGATATCCATCGCTACTACTTCTTCTTTCG |
| | Gbs-LAZY3-pGEX4T-1-F | CGCGTGGATCCCCGGAATTCATGAAGATTTTTAGTTGGGTTCAAAGAAAGCT |
| | Gbs-LAZY3-pGEX4T-1-R | TCACGATGCGGCCGCTCGAGGTTGGCGTCAACACTTGAACG |
| | Gbs-LAZY4-pGEX4T-1-F | CGCGTGGATCCCCGGAATTCATGAAGTTTTTCGGGTGGATG |
| | Gbs-LAZY4-pGEX4T-1-R | TCACGATGCGGCCGCTCGAGTCACCCCCCATCGTTACTGCTTCG |

| | | |
|---|---|---|
| | LAZY4-BH1-mut-F | GCGCTCTTGACGGCGGCGAGTGCGGCGGCGGCGTCTGATGTGAATCGAGAA |
| | LAZY4-BH1-mut-R | CGCCGCCGCCGCACTCGCCGCCGTCAAGAGCGCCGTCAGCTCCTTCTGTAG |
| | LAZY4-BH2-mut-F | GCGAACGCGACGGCGGCGAATGCGGCGGATTTGAGCGCGACCTCTGTTTCTCATCTTCTCGCGGCGATGTTTGTCTGTACAGAAGGT |
| | LAZY4-BH2-mut-R | CGCCGCGAGAAGATGAGAAACAGAGGTCGCGCTCAAATCCGCCGCATTCGCCGCCGTCGCGTTCGCGCTCTCTGTAGAAATAGCTTT |
| | MKK5$^{DD}$-pET28a-F | GATCCCCGGAATTCGAATTCATGAAACCGATTCAATCTCCTTCT |
| | MKK5$^{DD}$-pET28a-R | TGCGGCCGCTCGAGCTCGAGAGAGGCAGAAGGAAGAGGACG |
| | MPK3-pET28a-F | GATCCCCGGAATTCGAATTCATGAACACCGGCGGTGGC |
| | MPK3-pET28a-R | TGCGGCCGCTCGAGCTCGAGACCGTATGTTGGATTGAGTGC |
| | MKK5$^{DD}$-pGEX4T-1-F | CGCGTGGATCCCCGGAATTCATGAAACCGATTCAATCTCC |
| | MKK5$^{DD}$-pGEX4T-1-R | TCACGATGCGGCCGCTCGAGCTAAGAGGCAGAAGGAAGAG |
| | MPK3-pGEX4T-1-F | CGCGTGGATCCCCGGAATTCATGAACACCGGCGG |
| | MPK3-pGEX4T-1-R | TCACGATGCGGCCGCTCGAGCTAACCGTATGTTGGATTGAG |
| pEC-Cas9 construct | rbcS-E9t_BamHI F | CGGGATCCAGAGCTTTCGTTCGTATCATCGG |
| | rbcS-E9t_EcoRI R | CGGAATTCGTTGTCAATCAATTGGCAAGTCA |
| | p35S-Cas9 modified F | CGAGGTCGACGGTATCGATGCTAGCA |
| | p35S-Cas9 modified R | AGCTTGCTAGCATCGATACCGTCGACCTCGGTAC |
| | EC1.2en EC1.1p_NheI F | CTAGCTAGCGAATAAAAGCATTTGCGTTTGGT |
| | EC1.2en EC1.1p_XhoI R | CCGCTCGAGTTCTCAACAGATTGATAAGGTCG |
